# Bias correction for integrated climate projection modeling and its effect on downstream biological models

**DOI:** 10.64898/2026.01.12.698884

**Authors:** Jennifer S. Bigman, Kelly A. Kearney, Kirstin K. Holsman

## Abstract

Projections of future conditions from Earth systems models (ESMs) are necessary to understand and predict effects of changing environmental conditions on biological systems. Such projections suffer from biases, or mismatches between model output and observations. While adjusting or bias-correcting model output is common, many methods exist with little understanding of their effects on forecasts of biological change. Here, we explore the bias-correction process and its effects on downstream predictive biological models. As an example, we use the Bering 10K, a downscaled ESM for a productive and economically important subarctic ecosystem. We first characterize existing biases for three categories of variables exhibiting different scales and challenges: bottom temperature, sea ice, and net primary production. We then apply eight bias-correction approaches to six indices generated from the three categories and quantify sources of uncertainty in the trajectories of these ecosystem variables. Finally, we demonstrate how different bias-correction approaches affect downstream biological models using three case studies: 1) fish thermal spawning habitat suitability, (2) predicted zooplankton abundance, and (3) match-mismatch of phytoplankton and zooplankton bloom timing. We find that biases manifest in absolute values over time and in the timing of seasonal events. Time series of all six indices differed depending on bias-correction method, differences that were propagated to downstream biological models. For a given year and simulation, depending on method, thermal spawning habitat suitability and zooplankton abundance differed up to 149% and 151%, and match-mismatch increased or did not change. Our work highlights that bias correction reduces mismatches between observations and model output but choosing an approach requires careful consideration as to not amplify and propagate bias in downstream biological models. To that end, we identify best practices for bias correcting global or regional ESMs, including a decision tree to help improve forecasts of the effects of climate change on biological systems.

## 1 Introduction

Earth system models (ESMs) are important tools in climate research as they produce realistic projections of future conditions under a range of greenhouse gas emission scenarios (Eyring et al. 2016, Kearney et al. 2021). ESMs have a range of applications, from informing mechanistic linkages among biogeochemical factors to predicting the effects of climate on various sectors, including public health, agriculture, and more generally, food security (Teutschbein & Seibert 2012, Hawkins et al. 2013, Vicedo-Cabrera et al, 2019). Increasingly, ESMs are being used in the management of living marine resources, particularly in the context of fisheries (Gaines et al. 2018, Free et al. 2020). For example, ESMs are used to predict how commercially important species’ distributions, size, and ultimately, productivity, will fare in the future (Cheung et al. 2013, Clarke et al. 2021, Holsman et al. 2020).

ESMs, like all models, are imperfect depictions of the real world, and as such, their representations of physical and biogeochemical variables differ from reality (Ho et al. 2012, Hempel et al. 2013, Maraun et al., 2017). Biases in ESMs arise for a number of reasons, including limited spatial resolution, imperfect numerical discretization schemes, incomplete knowledge of the physical and biogeochemical interactions in the system, and the need to parameterize processes in terms of explicitly solved/modeled fields (Ho et al. 2012, Drenkard et al. 2021, Dinh & Aires 2023). Biases can manifest as offsets in the mean value of a variable of interest, as well as in the spatial and temporal characteristics of a variable, and can vary depending on the spatial and temporal scales of interest (Ho et al. 2012, Dinh & Aires 2023).

Model biases introduce a number of difficulties when attempting to use ESM output to investigate and predict ecosystem changes (Teutschbein & Seibert 2012, Maio et al. 2016). The growth, recruitment, and survival of many species are often tied to specific environmental conditions with particular spatial and temporal characteristics (Grüss et al. 2021, Morrongiello et al. 2021, Bigman et al. *in review*). Even small biases can lead to simulated historical conditions that cannot be directly reconciled with observations, thereby making interpretation of projected future changes challenging (Drenkard et al. 2021, Dinh & Aires 2023, Pozo Buil et al. 2023). Thus, it has become typical to apply some type of calibration, or bias correction, to ESM output to deal with these biases before forcing a downstream biological model.

Bias correction for ESMs to terrestrial applications have been well-studied, with much discussion on the motivation for bias correction and the risks of its misapplication (Ho et al. 2012; Maio et al 2016, Dinh & Aires 2023). However, there remains little consensus regarding which of the many methods to apply, exactly how and when to apply them, how marine applications may differ, and, importantly, how they may affect the outcome of downstream models that aim to forecast future biological change (Drenkard et al. 2021, Pozo Buil et al. 2023). The vast majority of the literature on this topic focuses primarily on a few limited variables (temperature and precipitation), often for terrestrial systems. These examples may not be representative of the full range of challenges encountered when attempting to adjust ESM output for the purpose of forcing biological models (Hawkins et al. 2013, Maio et al. 2016).

Modeling frameworks designed to investigate the long-term impact of climate change on aspects of biological systems (e.g., fishes) often involve a chain of one-way models. ESM output can be used directly as input to biological or ecological models (hereafter, we call these ‘downstream biological models,’ which encompasses a wide variety of biological and ecological modeling methodologies, including food web models, single and multispecies stock assessment models, species distribution models, etc. that use climate model output as input; e.g., in Cheung et al. 2013, Clarke et al. 2021, Bigman et al. 2023). Alternatively, the output from a global ESM can be used to force a regional ocean model, which acts to dynamically downscale the global ESM, with the regional model output then being used to drive downstream biological models (Drenkard et al. 2021, Pozo Buil et a. 2023). Regional ocean models are typically used in this context to ameliorate biases of global or parent ESMs, particularly those introduced by coarser spatial scale or less accurate representation of local processes of these models. However, regional models can exacerbate other biases inherited from the parent ESM or introduce new biases based on their own imperfect representation of certain processes (Drenkard et al. 2021).

Here, we focus on bias correction of a dynamically downscaled model, though our analysis is also applicable to direct use of ESM output for biological and ecological modeling. Specifically, we examine how different bias-correction methods affect the trajectories of abiotic and biotic variables from downscaled ESMs and then explore how these differences are propagated to downstream biological models. To do so, we use the Bering 10K Regional Ocean Model (hereafter, ‘Bering10K’) to explore the bias correction process. The Bering 10K is a high-resolution ocean, sea ice, and biogeochemical model that has been used to downscale a suite of ESM-derived abiotic and biotic variables for the Bering Sea and northern Gulf of Alaska (Kearney et al. 2020, Cheng et al. 2021, Hermann et al. 2021). The Bering10K output is used to drive fisheries and other biological models and predict how many biological aspects relevant to fisheries management will fare in the future (and is part of the Alaska Climate Integrated Modeling [ACLIM] project; see Hollowed et al. 2020, Holsman et al. 2020 for more information). First, we characterize the patterns of bias (mismatch) seen in three variables — bottom temperature, sea ice, and primary production — that are representative of different bias-related challenges. From these variables, we build a set of indices representative of those commonly used to force downstream biological models, apply a variety of bias correction methodologies to these selected indices, and characterize the uncertainty that arises in these time series due to bias correction choices. We then use three case studies to illustrate how different bias-correction methods affect the outcome of downstream biological models: forecasting 1) spawning habitat for Pacific cod based on bottom temperature, (2) zooplankton abundance based on sea ice, and 3) match-mismatch of phytoplankton and zooplankton bloom timing. Finally, we synthesize these steps to identify best practices for bias correction, as well as present a decision tree to help those using ESMs in downstream biological models navigate the bias correction process.

## 2 Methods

### 2.1 The Bering10K model simulations

Bering10K model is an instance of the Regional Ocean Modeling System (ROMS), with a domain that spans the Bering Sea and northern Gulf of Alaska. It includes explicit ocean, sea ice, and biogeochemical components (Hollowed et al., 2020, Kearney et al. 2020, Cheng et al. 2021, Hermann et al. 2021). The output we use in this study spans several simulations: the hindcast and a suite of downscaled long-term forecast simulations (Figure 1).

**Figure 1:**
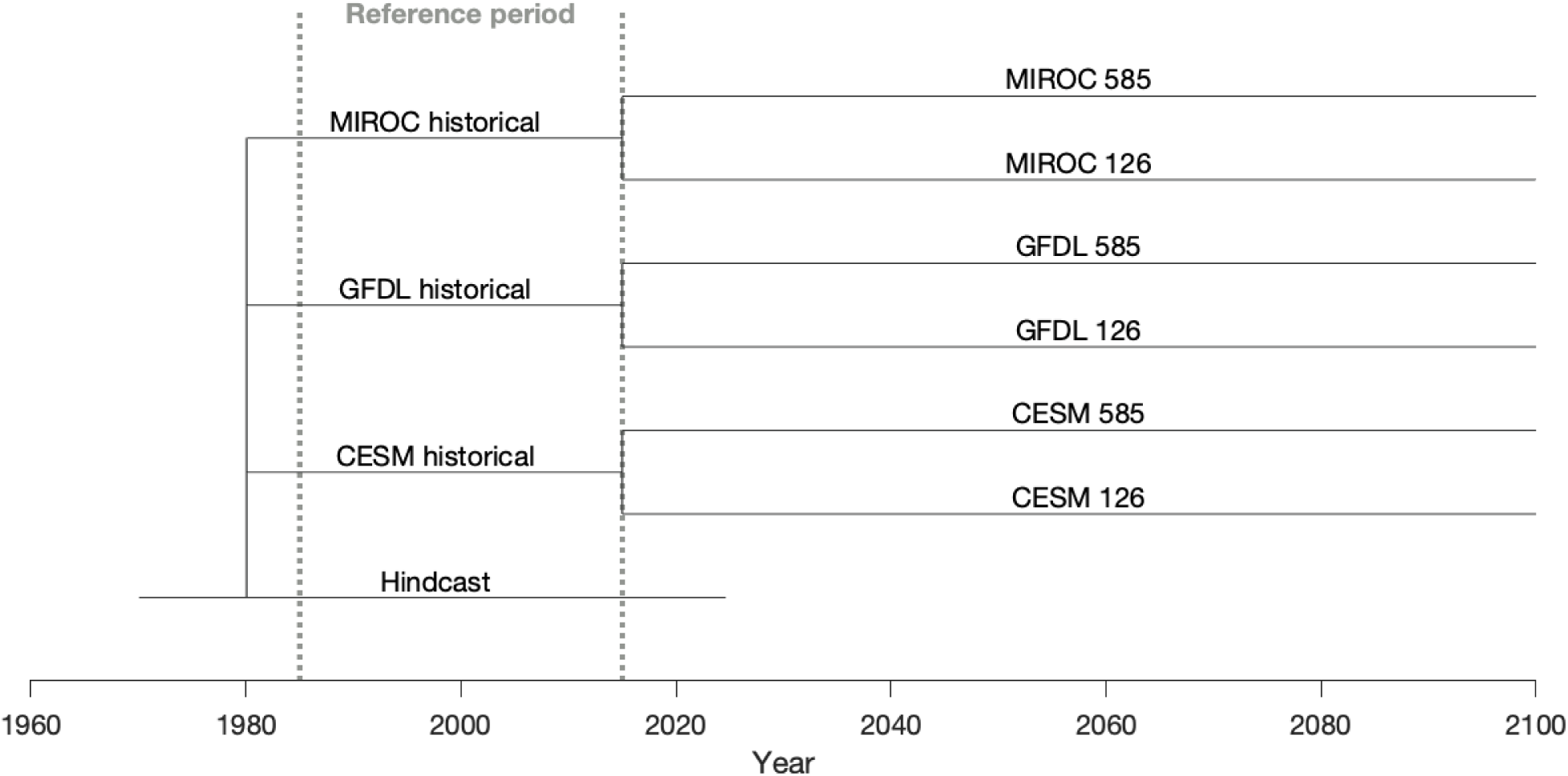
Schematic of simulation timelines. The historical ESM simulations are initialized from the hindcast simulation in January 1980. The ESM Shared Socioeconomic Pathways (SSP) simulations then initialize from their respective historical simulations at the start of 2015.

The hindcast simulation is driven by surface and boundary conditions from a variety of reanalysis datasets: the Coordinated Ocean Reference Experiment v2 (COREv2, 1970-1994, Large and Yeager 2009), Climate Forecast System Reanalysis (CFSR, 1995-2011, Saha et al. 2010), and Climate Forecast System Operational Analysis (COREv2, 2011-present, Saha et al. 2014). The hindcast simulation has been extensively documented previously and demonstrates skill in capturing real-world variations in key physical and biological variables (Kearney et al. 2020, Kearney 2021). In this paper (and throughout the ACLIM project), the hindcast simulation is used as our observational dataset.

The long-term forecast simulations are downscaled from selected parent models of the Coupled Model Intercomparison Project Phase 6 (CMIP 6): the Community Earth System Model version 2 with Community Atmospheric Model version 6 (CESM2-CAM6, hereafter ‘CESM’), Geophysical Fluid Dynamics Laboratory Earth System Model version 4.1 (GFDL-ESM4; hereafter, ‘GFDL’), and Model for Interdisciplinary Research on Climate-Earth System Version 2 for Long-term simulations (MIROC-ES2L; hereafter, ‘MIROC’). The latter portion of the historical simulation (1985 - 2015) and two different emission scenarios (2015-2099) — low (SSP126) and high (SSP585) — were downscaled for each parent model. These specific ESMs and emission scenarios were chosen to capture as much of the envelope of uncertainty from the larger CMIP6 suite as possible given the computing constraints that prohibited downscaling the entire suite.

To clarify terminology, throughout this paper we use the term “parent model” to refer to the ESM or reanalysis model (hindcast, GFDL, CESM, or MIROC) used as input to the Bering10K regional model, and “simulation” to refer to a specific run of the Bering10K model. Our suite includes a single hindcast simulation and two projection simulations per ESM. Each pair of projection simulations share the same forcings during the historical period and then split into the different emissions scenarios.

### 2.2 Bias correction of Bering10K output

#### 2.2.1 Why, when, and how to bias correct

The first consideration related to bias correction is whether to bias correct at all. Given the lack of consensus regarding bias correction best practices (Drenkard et al. 2021, Dinh & Aires et al. 2023, Pozo Buil et al. 2023) and the potential for various methods to be misapplied or misinterpreted (Maraun et al., 2017), it can be argued that bias correction should be avoided entirely (Ehret et al., 2012). However, we found that practical interpretation of projected change was quite limited when bias correction was avoided. Many species have narrow tolerances for a range of environmental conditions, as well as thresholds and/or tipping points, such that even small biases can lead to significant misrepresentation of their ecological dynamics.

Given the need to adjust for these biases, one of the first considerations when bias correcting within a dynamic downscaling framework is where in the process to apply bias correction. Bias correction in dynamical downscaling frameworks is usually applied either to the output from the ESMs that is used as input forcing datasets for the regional model (e.g., before dynamical downscaling; Pozo Buil et al. 2023) or to the output variables produced by the regional model after it has been downscaled (e.g., Bigman et al. 2023, Holsman et al. 2020). Importantly, biophysical models at both global and regional scales have a high degree of nonlinearity in their governing equations and thus, bias correcting the inputs to these models (prior to downscaling) will not yield the same result as bias correcting the output. The Bering10K was not bias corrected prior to downscaling for a few reasons. First, the Bering Sea region is strongly influenced by seasonal sea ice dynamics (Stabeno et al. 2012). Sea ice is modulated by a variety of atmospheric forcing elements, including the local air temperature and wind patterns, which are themselves shaped by the local radiative balance, air pressure, etc. These variables are all provided individually as atmospheric forcing to the regional ocean model (Kearney et al. 2020, Cheng et al. 2021, Hermann et al. 2021). If one were to bias correct each forcing variable independently, it could easily decouple important relationships between each variable, creating forcing that appeared correct but at the cost of erasing potentially important feedbacks between larger-scale processes. Second, the ESM-forced Bering10K simulations are used in conjunction with a variety of different downstream ecological models. Application of bias correction to model output post downscaling provided a more flexible framework to tailor to individual applications, given that it does not require rerunning the computationally expensive regional model.

#### 2.2.2 Characterizing model bias for climate indices

We selected three output variables from our regional model simulations as the base for our set of indices: bottom temperature, phytoplankton production, and fraction ice coverage. These three variables were selected to be representative of the behavior and challenges associated with the wider suite of ocean, ice, and biogeochemical model variables that can be used to build ecologically-relevant indices.

Bottom temperature is — like other variants of ocean temperature — widely used as a biological indicator. In the Bering Sea shelf region, variability in bottom temperature includes a seasonal cycle as well as interannual variability in mean; the latter is often characterized by warm and cold stanzas of one to four years in length (Stabeno et al. 2012). Superimposed on this seasonal and interannual variability is a clear long-term trend in the mean associated with anthropogenic climate change (Kearney et al. 2020, Cheng et al. 2021, Hermann et al. 2021). At the regional scale, biases in bottom temperature present primarily as offsets in the mean and over- or underestimation of the amplitude of the seasonal cycle (Kearney et al. 2020, Cheng et al. 2021, Hermann et al. 2021).

Net phytoplankton production is representative of plankton growth rate and biomass variables; it is characterized primarily by a strong seasonal cycle with near-zero values during the winter months, a strong spring bloom peak, and sometimes a smaller peak in the late summer or early fall (Kearney et al. 2020, Cheng et al. 2021). Interannual variability includes variations in the amplitude and timing of these peaks during the spring-to-fall period, with little to no variability during the winter period. Long-term climate change signals manifest as changes in peak timing, height, and prominence, but without the clear directional agreement across parent models that is seen in temperature. Regional biases are also characterized by mismatches in the timing, height, and prominence of these peaks (Kearney et al. 2020, Cheng et al. 2021). Plankton growth and mortality processes are nonlinear functions of physical variables including temperature, nutrients and light availability, and therefore biases in production reflect a combination of the biases of the parent forcing model’s physical drivers (e.g., the ESM temperature bias) and of the regional model itself (e.g. how production responds to temperature). Due to the timescale of the processes involved and the geometry of the Bering10K domain, biases in ESM phytoplankton production provided through initial and boundary conditions do not propagate to the Bering shelf region where our indices were calculated.

The final variable we focus on, fractional sea ice coverage, is also characterized by a strong seasonal cycle, with values ranging from zero during the summer months to near one (total coverage) during peak winter months. Interannual variability is primarily characterized by variations in the peak value, with little to no variation during the zero-ice summer months (Cheng et al. 2021, Hermann et al. 2021). In the Bering10K, the distribution of this variable is constrained between zero and one. The Bering10K model is initialized with no ice and uses primarily closed ice boundary conditions, so, like production, the biases in this variable are primarily a function of biases in other physical drivers like temperature rather than being inherited from the parent model boundary forcing (Kearney et al. 2020, Cheng et al. 2021, Hermann et al. 2021). Biases manifest primarily as offsets in maximum peak value along with small offsets in the timing of sea ice advance and retreat. The climate change signal in this variable is reflected in a decrease in peak ice coverage, but because of the constraints on value to stay between zero and one, the trend is not as near-linear as seen in temperature itself.

Each of the three base output variables were archived at a weekly time step across all simulations. We first extracted one-dimensional weekly time series for each simulation by extracting and spatially-averaging values across the southeastern Bering Sea shelf as defined by NOAA Alaska Fisheries Science Center’s bottom trawl groundfish management strata polygons (Bakkala et al. 1985). Starting with these three variables, we calculated six different annually-resolved biological indices (as the annual time step is commonly used to force biological models): annually-averaged bottom temperature, annually-averaged phytoplankton production, spring bloom start date, monthly-averaged September phytoplankton production, maximum fraction ice coverage, and number of ice-free weeks per year. These were chosen to reflect how different bias correction methods may impact indices that reflect different aspects of a time series (central value vs. extrema, phenology, and proximity to key thresholds).

#### 2.2.3 Bias correction methods

The most widely used bias correction methods are based on the relationship between modeled variables, both during a present-day reference period and a future projected period, and a best estimate of the real-world state of those variables during the present-day period (Ho et al. 2012, Maio et al. 2016). This latter dataset is often derived from a gridded observation-based data product, data-assimilating model reanalysis, or a reanalysis-forced hindcast simulation; we will refer to this as the observed dataset or ‘observations’ for shorthand. Here, we applied several different bias-correction methods to the above weekly and/or annual time series that are based on this general approach (Figure 2). Each method adjusted raw values from the simulation (*x*) by comparing the distributions of the historical portions of the ESM projection simulations to the hindcast simulation during an overlapping reference period of 1985–2015 and remapping to a bias-corrected time series (*x*′). This 30-year reference period was chosen to encompass typical decadal variability in the region while avoiding spinup effects associated with initialization of the downscaled ESM models. The bias correction methods were applied as follows:

1. No bias correction: Annual indices were calculated directly from the unadjusted weekly time series.
2. Bias removal and rescaling: This method has been called the ‘delta’ method (with unequal variance, see Holsman et al. 2020) and follows the Ho et al. (2012) semiparametric approach to bias correction:

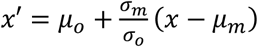

We note that this method assumes values in both the observational and model datasets over the reference period can be described by empirical distributions defined in terms of location (*μ*) and scale of variance (*σ*) parameters. In most applications, including in this one, *μ* and *σ* are set to the mean and standard deviation, respectively, of the datasets; this appropriately describes the distributions if they are normally distributed but may be a suboptimal simplification otherwise. In our application, *μ*_*o*_ and *μ*_*m*_ are the mean of observed (hindcast) and modeled (historical portion of the forecast simulation) time series, respectively, at weekly resolution from 1985–2015; *σ*_*o*_ and *σ*_*m*_ are the standard deviations of the same weekly data. Annual indices were calculated from the bias-corrected weekly data.
3. Bias removal, no rescaling: This method has also been called the ‘delta’ method (but assumes equal variance) and is identical to method #2 but without the 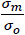 scaling factor. This reduces to a simple removal of mean bias.
4. Climatological bias removal and rescaling: This variant on method #2 uses the same equation but calculates *μ*_*o*_, *μ*_*m*_, *σ*_*o*_, and *σ*_*m*_ separately for each week of the year across the reference period. This method attempts to account for the differing biases and scaling factors during different seasons. In common practice, this sort of climatological bias correction is typically applied at a monthly or seasonal (quarterly) resolution; we instead apply it at a weekly resolution to highlight artifacts that can arise due to phenological biases that are exacerbated with finer temporal resolution of the climatological reference values.
5. Climatological bias removal, no rescaling: This method is identical to method #4 but without the 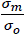 scaling factor.
6. Quantile delta mapping, multiplicative: This method follows that proposed by Cannon et al. (2015). This method first remaps values in the model reference period to observations using the empirical cumulative distribution functions (CDFs) for each; e.g., a value that corresponds to the n^th^ percentile of the modeled reference period would be transformed to the value matching the n^th^ percentile of the observations. For future periods, CDFs are also calculated for each point using a sliding window of the same width as the reference period, and the relative delta (n^th^ percentile of future period divided by n^th^ percentile of reference period) is applied as a scaling factor to the n^th^ percentile of the observations to create the final bias-corrected projection.
7. Quantile delta mapping, additive: Similar to method #6, this follows the Cannon et al. (2015) remapping process but preserves the absolute change rather than the relative change, such that a delta offset value (n^th^ percentile of future period minus n^th^ percentile of reference period) is added to the n^th^ percentile of the observations.
8. Bias removal and rescaling, post-annual: This method is identical to method #2, except that the annual indices are calculated from the un-bias-corrected weekly time series first, and then bias correction is applied to the resulting annual time series.
9. Bias removal, no rescaling, post-annual: This method is identical to method #8 but without the 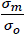 scaling factor.

We made a few adjustments to these equations to deal with numerical discontinuities across these methods. First, when applying any variant with scaling method, we capped the absolute value of the scale factor at 10. This cap was chosen to avoid very large absolute scale factors resulting when standard deviation was near zero in the reference model, under the assumption that variability between models would not vary by more than an order of magnitude except when that variability approaches zero. Discontinuities also appeared during our first attempt at applying the scaled quantile delta mapping algorithms to bottom temperature. These were resolved by converting from °C to °K to avoid numerical discontinuities that would otherwise occur when temperature approached 0°C (a numerical threshold that does not have a corresponding physical meaning in saltwater). However, for ease of interpretation, we present all results in °C.

**Figure 2.**
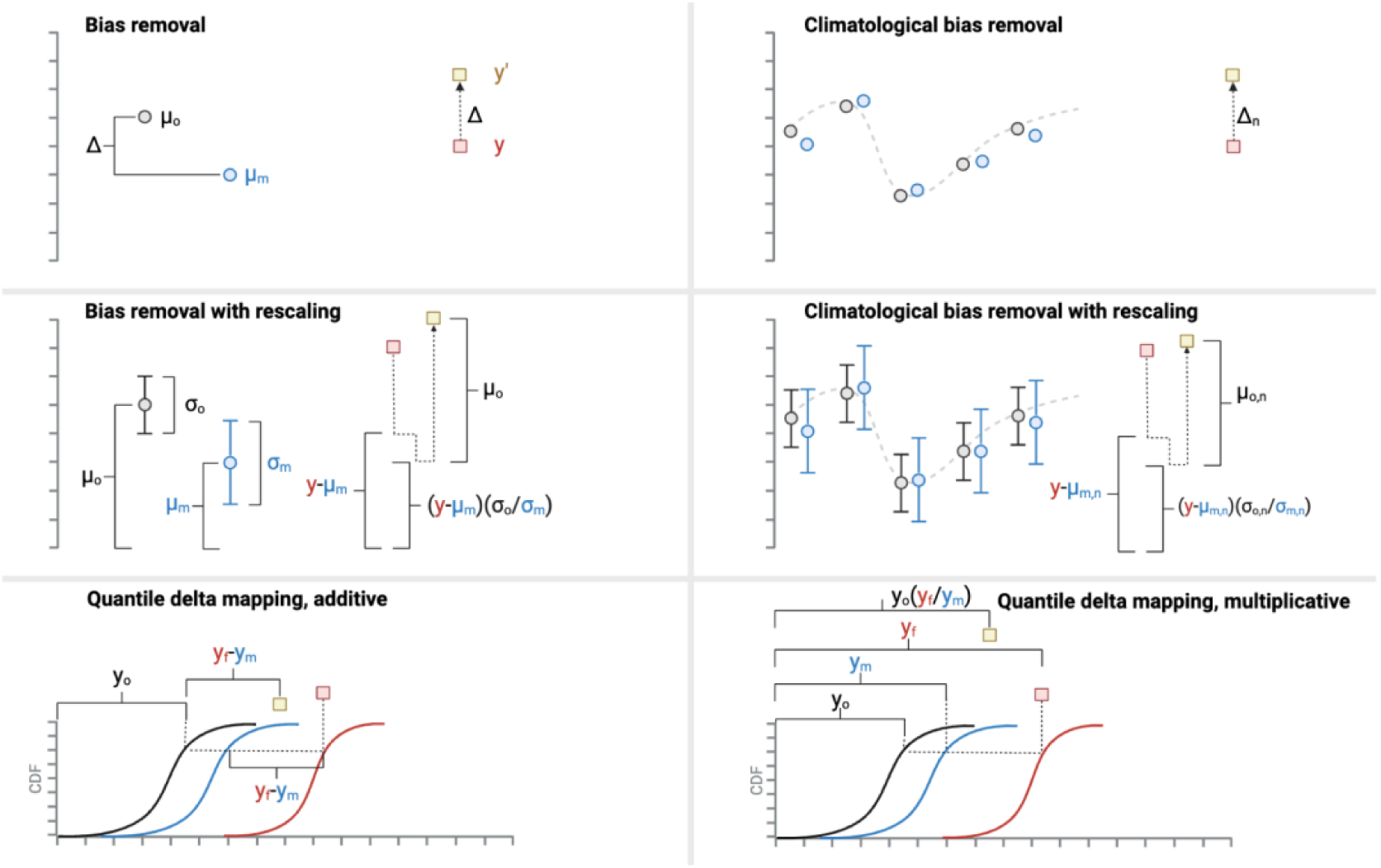
Six of the eight bias-correction approaches applied in this study. The other two, post-annual bias removal with and without rescaling are similar to bias removal and bias removal with rescaling, but before any bias correction, are averaged by year, generating annual indices, and then bias corrected. The x-axis represents time (e.g., year) and the y-axis represents the scale of the variable of interest.

#### 2.2.4 Bias uncertainty relative to other sources of projection uncertainty

The CMIP6 ScenarioMIP experiment, and by extension the ACLIM experimental setup, was designed to encompass many different sources of uncertainty that could contribute to end-of-century projections (O’Neill et al., 2016, Hollowed et al., 2020). These uncertainties can be high, and previous studies have classified the sources into three primary categories: internal variability uncertainty, model uncertainty, and scenario uncertainty (Figure 3; Hawkins and Sutton 2009, 2011, Frölicher et al. 2016). Internal variability uncertainty (‘internal variability’) can be defined as the natural decadal- and interannual-scale variability in the system, which is distinguished from external forcing, e.g., due to anthropogenic climate change (Deser et al. 2009, 2011). For example, internal variability could reflect climate oscillations like the El Niño-Southern Oscillation (ENSO) and is usually the dominant source of uncertainty over shorter (decadal) time frames (Hawkins and Sutton 2011, Frölicher et al. 2016). Model uncertainty (also called structural uncertainty) encompasses uncertainty deriving from our imperfect understanding of processes and/or our ability to numerically represent them in models. This is typically quantified by comparing the range of simulation output across different ESMs using the same or similar external forcing. (Hawkins and Sutton 2009, 2011, Frölicher et al. 2016). Such differences may arise from, for example, the use of different physical and biogeochemical equations, parameters, and numerical schemes. Scenario uncertainty reflects the range of potential alternative futures depending on policy decisions and technological or sociological considerations (Hawkins and Sutton 2009, Frölicher et al. 2016). Within CMIP6, this uncertainty is encapsulated by the various Shared Socioeconomic Pathways (SSPs) and their accompanying emissions scenarios.

**Figure 3.**
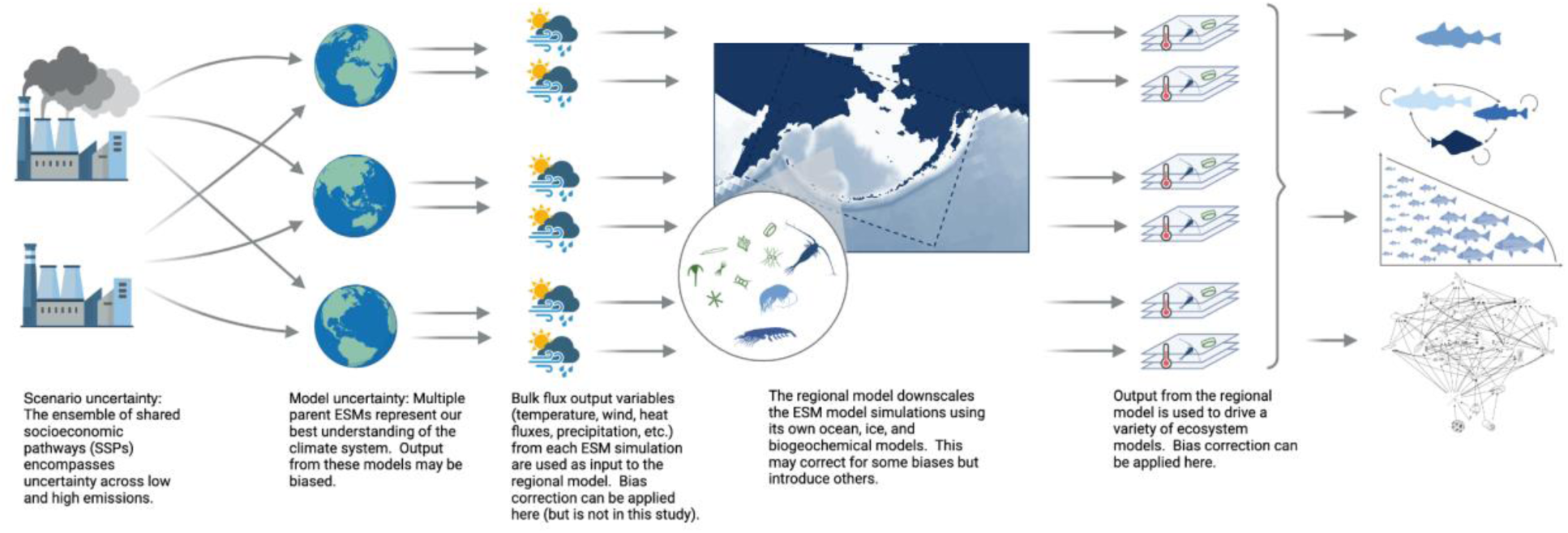
A conceptual diagram of the types of uncertainty in the CMIP6 suite of models, as well as when bias correction could occur.

To disentangle sources of uncertainty in the suite of Bering 10K simulations and to understand how much uncertainty bias correction introduced, we quantified the above three sources of uncertainty for all six indices following Frölicher et al. (2016). We also calculated a new measure of uncertainty - ‘bias-correction uncertainty,’ which measures how much uncertainty arises due to different bias correction methods (Figure 3). Internal variability was calculated as the multimodel mean of one standard deviation of the 30-year historical period (1985-2015) of each simulation. We assume (following Frölicher et al. 2016) that this uncertainty is constant through time. Model uncertainty for each year was calculated by first applying a 10-year moving mean to each index time series and then quantifying the mean of the (one) standard deviation of the individual low emission scenario simulation across parent models (e.g., standard deviation of CESM SSP126, MIROC SSP126, and GFDL SSP126) and the (one) standard deviation of the high emission scenario simulations (e.g., standard deviation of CESM SSP585, MIROC SSP585, and GFDL SSP585). Finally, the scenario uncertainty for each year is the difference between the multimodel mean of the high emission scenario simulations (e.g., mean of CESM SSP585, MIROC SSP585, and GFDL SSP585) and the multimodel mean of the low emission scenario simulations (e.g., mean of CESM SSP126, MIROC SSP126, and GFDL SSP126), again calculated after application of a 10-year moving mean. The 10-year moving mean is intended to minimize the contribution of internal variability to either model and scenario uncertainty (Frölicher et al. 2016). The novel source of uncertainty calculated here - the bias-correction uncertainty - was calculated as the multimodel and multiscenario mean of (one) standard deviation across bias correction methods.

### 2.3 Biological model case studies

#### 2.3.1 Forecasting Pacific cod thermal spawning habitat

Spawning dynamics are tightly tied to thermal conditions for many groundfish species, leading many to use hindcasts and forecasts of temperature to predict how the availability of spawning habitat may change over time as waters warm (Dahlke et al. 2018, Cote et al. 2021, Bigman et al. 2023). This is especially true for Pacific cod, as this species lays demersal eggs that have a narrow thermal range that confers successful development and hatching (Alderdice & Forrester 1971, Laurel & Rogers 2020). Here, we use this tight connection between temperature and the thermal response of embryos to compare the effect of bias correction methods on forecasts of thermal spawning habitat (i.e., based on temperature). Specifically, we calculated an annual index of Pacific cod thermal spawning habitat suitability by coupling nine time series of bottom temperature (raw and those resulting from eight bias-correction approaches) with an established experimentally-derived relationship between hatch success and temperature (see Laurel & Rogers 2020, Bigman et al. 2023). This resulted in a time series of the proportion of successful hatch for each year, bias correction approach, and simulation (e.g., low emission scenario of CESM). We then compared this index across all time series to understand how the magnitude and trend of the index of thermal spawning habitat suitability differed among bias correction methods.

#### 2.3.2 Sea ice & zooplankton

Zooplankton abundance in the Bering Sea is known to be closely related to sea ice such that reduced sea ice is correlated with lower zooplankton abundance (Duffy-Anderson et al. 2019, Kimmel et al. 2023). We used this relationship to understand how the different ways of bias correcting sea ice time series from the Bering10K affected predictions of zooplankton abundance. To do so, we obtained annual zooplankton abundance (number per meter^3^ of large copepods [> 2mm]) from multidisciplinary ecosystem surveys conducted in the spring and summer by the National Marine Fisheries Service Alaska Fisheries Science Center in Seattle, WA for all years available (1995, 1997-2000, 2003-2022). These surveys sample zooplankton via oblique tows of paired bongo nets (20 cm frame, 150 um mesh and a 60 cm frame, 333 um mesh; for more detail, please see Kimmel et al. 2018, Kimmel et al. 2024). We fit a mixed effects model with a gamma response distribution and log link to examine how observed zooplankton abundance varied with sea ice from the Bering10K. We included a random effect of year to account for other differences aside from those due to sea ice across years; we also examined whether a random effect of survey was needed, but it was not supported (i.e., AIC value was greater). Next, we used this model to predict zooplankton abundance from the nine projection time series of sea ice (raw plus eight bias correction approaches). We then compared these predicted time series of zooplankton abundance to understand how the different bias correction approaches affect forecasts of zooplankton abundance.

#### 2.3.3 Match-mismatch of phytoplankton bloom and peak zooplankton abundance

The spring phytoplankton bloom is a cornerstone of high-latitude systems and provides resources for a range of organisms, which then serve as prey for higher trophic levels (Ji et al. 2010, Nielsen et al. 2024). As environmental conditions change, it is expected that the timing of the spring phytoplankton bloom will shift but it is uncertain the effect this will have on other trophic levels and the ecosystems in which they live (Cushing 1990, Nielsen et al. 2024). For example, it is unclear whether and how much this will affect zooplankton populations, which consume phytoplankton and transfer energy to higher trophic levels. To understand how the match-mismatch of bloom timing for lower trophic levels may change in the future, we assessed whether the match - mismatch of the relative timings of the spring phytoplankton bloom and zooplankton peak abundance over time varied across the eight different bias correction time series and the raw time series. Specifically, we calculated the number of days between the peak phytoplankton bloom and the timing of peak zooplankton abundance. For each of the eight bias-corrected time series and the raw time series, and plotted how this value changed over time for the low and high emission scenarios. Within each emission scenario, the peak day of year for both phytoplankton and zooplankton were averaged (i.e., across the three parent models, CESM, MIROC, GFDL).

All model output, data, and code can be found here: (https://github.com/kakearney/supplementary-data-bias).

## 3 Results

### 3.1 Characterizing model bias for climate indices

The three parent ESMs used in this study were chosen to bracket the envelope of thermal sensitivity in the larger CMIP6 suite, and their bottom temperature biases reflect this spread. CESM is the warmest of the three ESMs; its historical-period seasonal amplitude is well-matched to the hindcast simulation but with a consistent bias of approximately 1°C year-round (Figure 4 top row). The GFDL and MIROC models show very similar mean historical seasonal cycles, both underestimating the seasonal amplitude with a cold bias that is stronger in summer than winter. Both seasonal and interannual variability in the MIROC model is the highest of the suite (Figure 4 top row).

**Figure 4.**
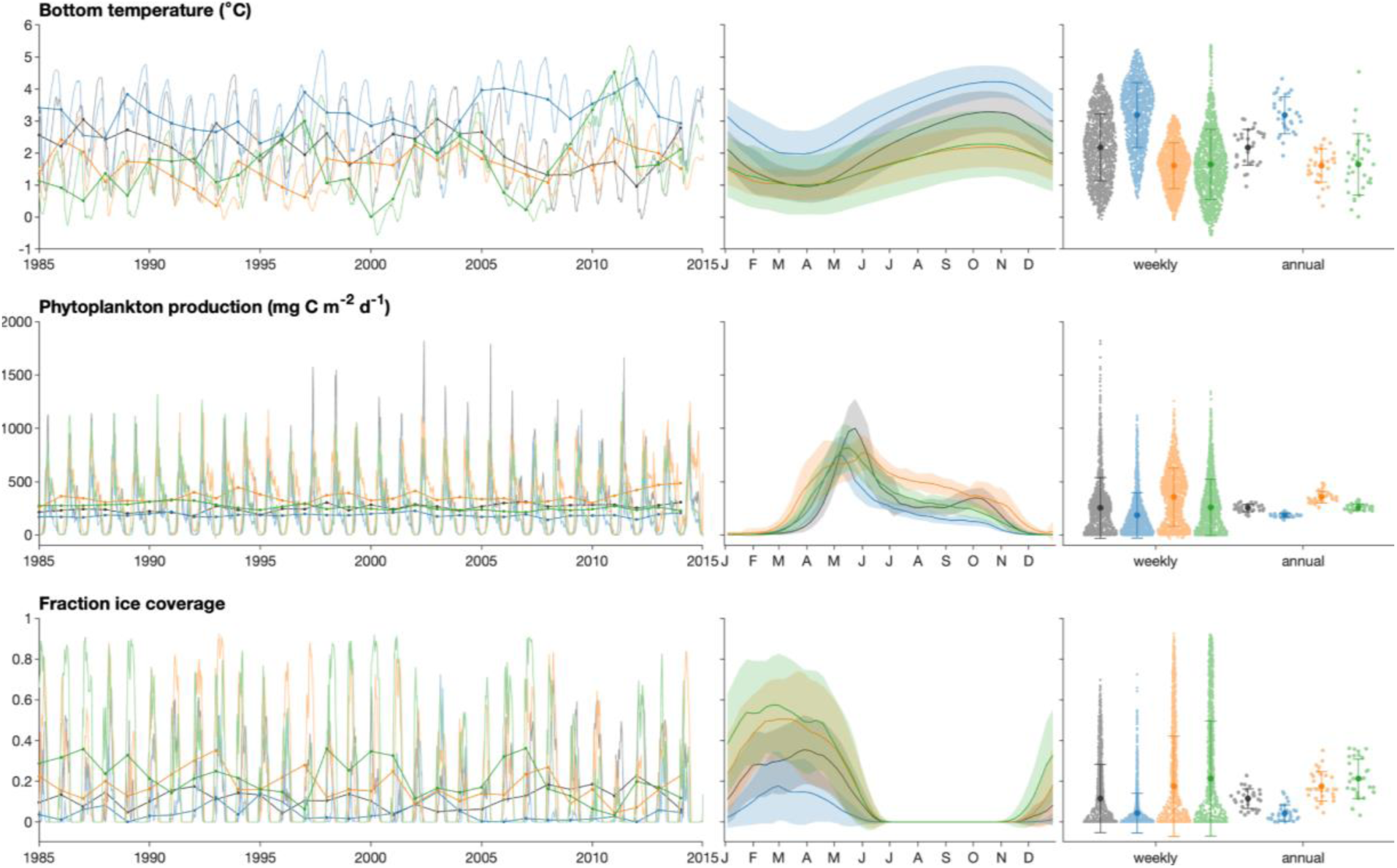
Left-most panels show the weekly (light colors) and annually-averaged (dark colors) time series for the three regional model output variables across the reference period. Colors indicate the hindcast (black), CESM (blue), GFDL (orange), and MIROC (green) models. The center panels show weekly climatological mean (lines) and standard deviation (shaded region) for each simulation across the same 30-year reference period. The right-most panels depict the mean and standard deviations (error bars) and approximate distribution (dots) of both the weekly and annually-averaged values across the 30-year reference period.

Net primary production biases are apparent in both the magnitude and timing of peak values (Figure 4 middle row). All three ESMs show an earlier onset of the spring bloom than in the hindcast. CESM and MIROC are both biased low throughout the bloom period and with early decay of net primary production; biases in magnitude and timing are stronger in CESM than MIROC. The GFDL model predicts a longer spring bloom than the hindcast, with an earlier rise and later decay of net primary production values and lower peak values. None of the three ESMs show a consistent presence of a fall bloom peak in the climatological mean, though fall blooms do appear in individual years throughout the reference period. The weekly distribution of values for this variable is bimodal with one peak in spring and one in fall (Figure 4, middle row). Higher distributions indicate the near-zero winter values and summer-to-fall intermediate values and a tail of lower-frequency high values correspond to spring. There is very little variability in the annual mean time series (the dark lines in the left panel of Figure 4 and the ‘annual’ values in the right column of Figure 4); the CESM model has a low bias at this timescale while the other two ESMs show close agreement with the hindcast simulation (Figure 4 middle row).

Biases in the fraction of sea ice coverage correspond closely to temperature biases (Figure 4 bottom row). The CESM model produces the least sea ice coverage, consistent with its warm temperature bias. It is the only model of the suite to simulate years with no sea ice on the southeastern Bering shelf during the reference period. Both the GFDL and MIROC models overestimate ice. The MIROC model has the highest interannual variability in peak ice. Timing of peak sea ice is similar across all models (Figure 4 bottom row).

### 3.2 Bias Correction Comparison

The various bias correction methods performed with varying effectiveness when applied to the three different (weekly) time series (Figure 5). All bias correction methods were able to correct for biases in the mean while preserving the seasonal cycle. However, we note the exaggeration of the seasonal cycle that is seen with the climatological bias removal and rescaling method (#5 in list in methods), which is particularly visible in the high emission CESM simulation (Figure 5 top panel). This artifact arises due to a relationship between bias and variability, especially apparent when bottom temperatures are near the freezing point.

**Figure 5.**
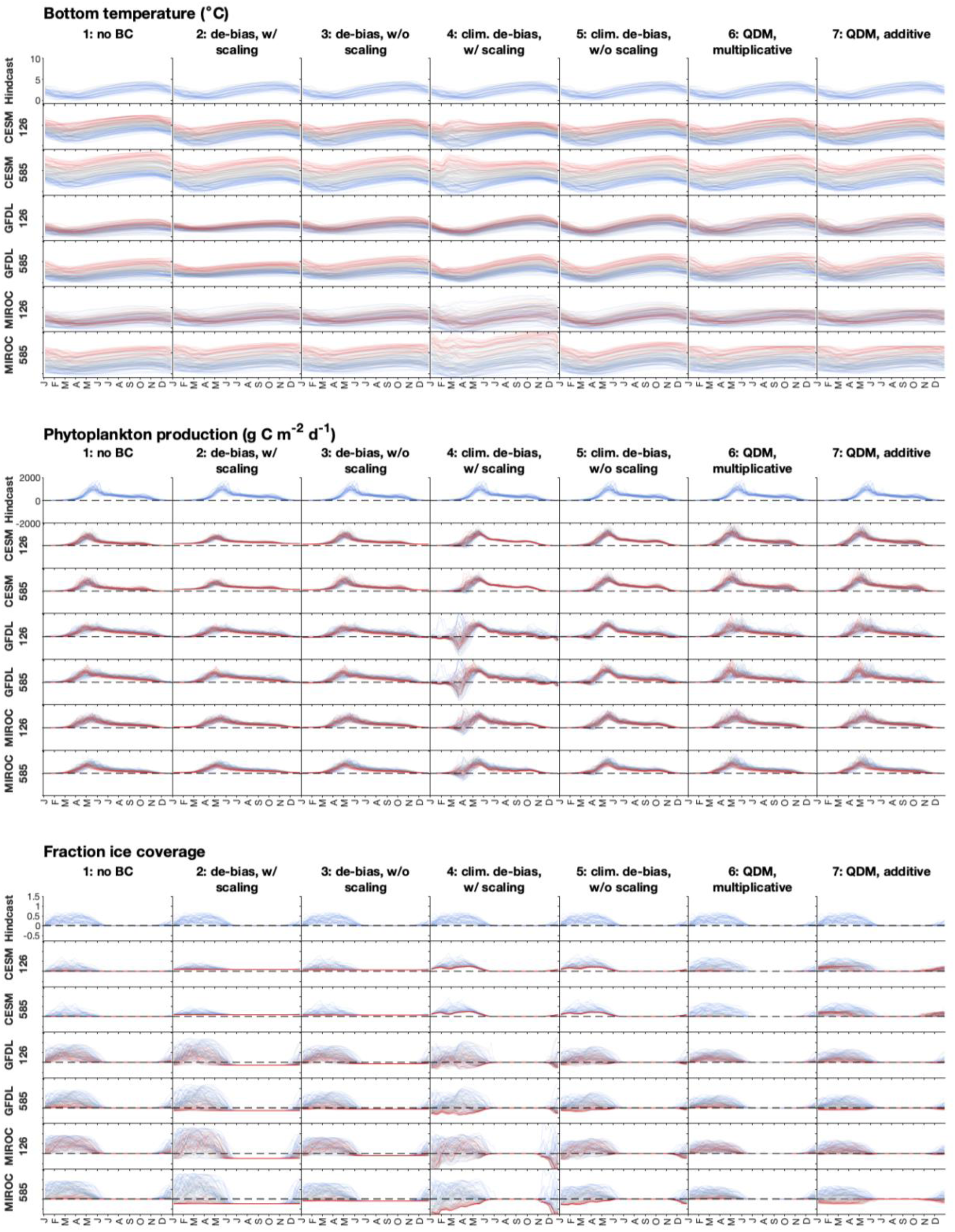
Bias-corrected time series vs. day of year for each simulation. Axes are arranged by parent model and SSP (rows) and bias correction method (column); all axes for each variable share the same vertical axis range. Blue lines indicate the 30 years from the reference period (1985–2015), red lines indicate the final 30 years of the long-term projections (2070–2100), and gray lines are the remaining years.

Bottom temperature showed the least variability across bias correction methods (Figure 5 top panel). Bottom temperature cannot drop below −1.7°C, and therefore, the models all display a decrease in variability correlated with lower mean temperatures. The hindcast, CESM, and GFDL models all appear to capture a similar underlying relationship between bottom temperature mean and interannual variability, with interannual variability highest during periods when the mean is approximately 2–3°C, and falling off to either side (Figure S1). The MIROC model sees a similar decrease in variability below 2°C, though its variability values are higher than the other models across the board. The climatological bias correction methods assume these variations in seasonal standard deviation are a function of fixed time, and when applied, this decouples the underlying relationship between mean and variability (Figure 5 top panel).

The bias correction methods showed a mix of issues when applied to net primary production (Figure 5 middle panel). The non-climatological bias correction methods (bias removal with and without rescaling, #2 and #3) both suffered from application of a constant offset across all seasons. As a result, models with a negative bias (CESM and MIROC) were consequently adjusted to always be above zero, even during the winter months when production should be negligible, while the positively-biased GFDL model was adjusted such that production dropped below zero during these months. The climatological methods (climatological bias removal with and without rescaling, #4 and #5) corrected for this constant offset issue. However, phenological mismatches between models led to a different problem, where the early onset of spring production was alternately adjusted downward in the pre-bloom period and upward toward the end of the bloom; the downward adjustments could lead to negative values in several of the models. In addition, as seen in the bottom temperature rescaled climatological application (climatological bias removal with rescaling, #6), variability is compressed when production is low due to the hard threshold on values at 0. The misalignment of these near-zero values across models lead to drastically scaled-up variability around the bloom onset period in the scaled version, exacerbating these overcorrections. The quantile delta mapping methods (#6 and #7) performed best for this variable, preserving temporal changes that occur in some simulations over time while maintaining near-zero but non-negative winter production values.

Sea ice fractional coverage, with its hard constraints on both the lower and upper end of its distribution, displayed many of the same issues as in the previous two variables, but to an even greater extent (Figure 5 bottom panel). Constant offset adjustments applied via the non-climatological bias removal methods (bias removal with and without rescaling, #2, and #3) led to obviously incorrect sea ice values during the summer months, with either year-round ice cover in the negatively-biased CESM model or frequent negative sea ice values in the positively-biased GFDL and MIROC models. The climatological bias correction variants (climatological bias removal with and without rescaling, #4 and #5) better maintained ice-free summer values but pushed values negative during the sea ice season for both the GFDL and MIROC variants, despite the equal scale assumption. For the bias correction methods with rescaling (methods #2, #4, and #8), values also often exceed one. The additive quantile delta mapping method (#7) similarly resulted in negative sea ice fraction toward the end of the century for positively-biased models when the absolute loss of sea ice surpassed the amount of sea ice available to lose from the downward-adjusted starting point. The multiplicative quantile delta mapping method (#6) was the only method that successfully constrained future values within the numerically-correct (0-1) range (Figure 5 bottom panel).

Many of the issues with variables crossing unrealistic physical or biological thresholds became hidden once converted to annual indices (#8 and #9 in methods list); all indices except maximum ice coverage stayed within their expected ranges on an annual scale (Figure 6). Mean bottom temperature annual indices clustered tightly across methods for both the GFDL and CESM models. However, the MIROC model’s overestimated variability led to exaggeration of the long-term trend with both the climatological and post-annual bias removal with rescaling method (#4 and #8). Likewise, net primary production clustered closely, but with exaggerated interannual variability in the GFDL model with the climatological bias removal no rescaling (#5) and post-annual bias removal methods (#8 and #9). The bloom start date index highlights the differences between methods in how they handle phenological indices. The climatological bias removal methods (#4 and 5) show high variability because of the dipole-like adjustments they apply in the vicinity of the bloom onset. The maximum ice coverage index highlights some of the issues that arise when applying the bias removal methods to a variable that is approaching a threshold. Opposite ends of this issue can be seen in the high emission CESM and MIROC simulations. Raw CESM, with its underestimated historical ice coverage, predicts near-zero sea ice by mid-century, but many of the bias-adjusted values plateau at non-zero values. The MIROC model, which overestimates historical period ice, predicts the most drastic decrease in ice across the long-term projection; many of the adjusted indices exaggerate this trend to the point of pushing values outside the feasible range (i.e., below 0 or above 1). The ice-free period index shows the highest cross-method spread. The CESM model alternatively predicts a future with either ever-present ice (bias removal without rescaling, #3) or almost nonexistent ice (post-annual bias removal and rescaling, #8), depending on the bias correction method. For the high-biased ice models, the multiplicative quantile delta method (#6) is an outlier to the other methods, predicting fewer ice-free weeks both during the historical period and in the future, and with a less steep long-term increase in ice-free weeks (Figure 6).

**Figure 6.**
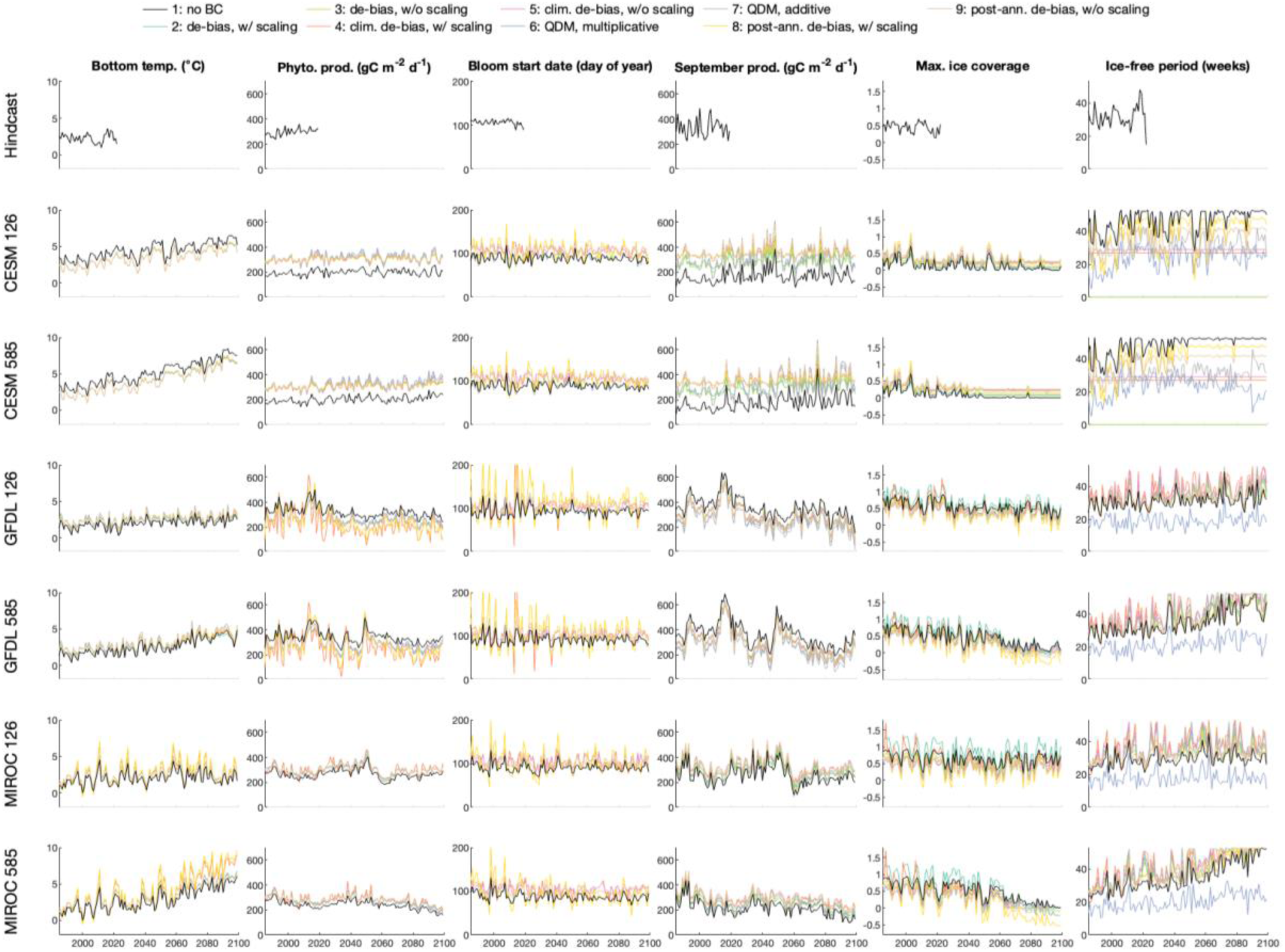
Annual time series for each index (columns) and simulation (parent model and SSP simulation; rows), colored by bias correction method.

#### Characterizing model uncertainty

We found that the relative contributions of the four sources of uncertainty - internal variability, model uncertainty, scenario uncertainty, and bias-correction uncertainty - differed depending on the category of variable (temperature, net primary production, sea ice) and which aspect of the time series they represent (e.g., metrics that characterize central tendency versus outliers or extrema, phenology, proximity to key thresholds; Figure 7). Model uncertainty was generally the largest source of uncertainty for annually-averaged bottom temperature, annually-averaged net primary production, and maximum fraction ice coverage. This type of uncertainty contributed at least 25% to the total for all six indices except for bloom start date, which was dominated by internal variability. For bloom start date, internal variability comprised at least half of the total uncertainty over the course of the 120-year time series. Internal variability was also an important source of uncertainty for monthly-averaged September net primary production, annual net primary production, and maximum fraction ice coverage, where this type of uncertainty comprised ∼ 25% or more for the duration of the time series (Figure 7).

**Figure 7.**
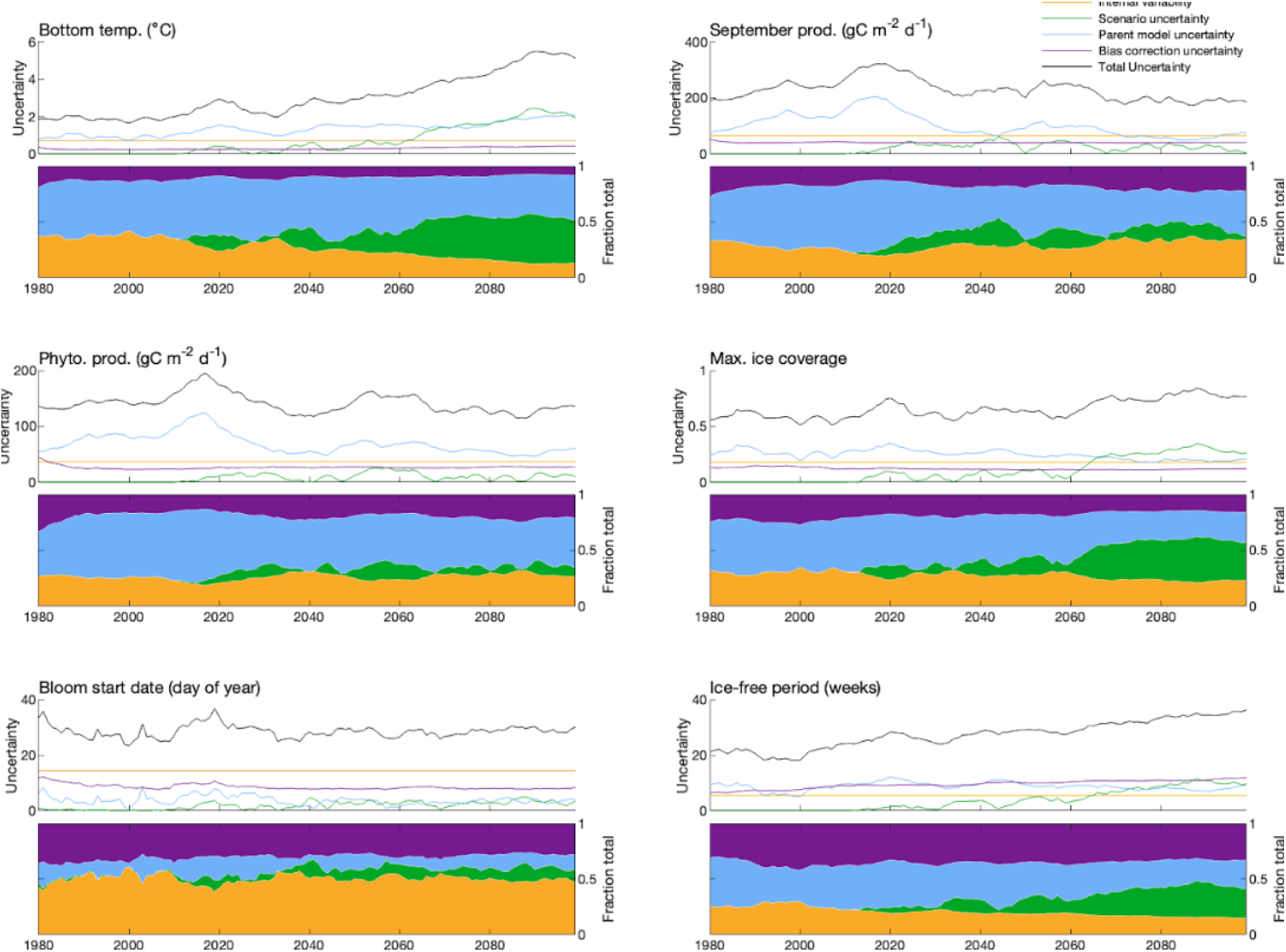
The uncertainty due to internal variability, differences across parent models and emission scenarios (scenario and parent model uncertainty), and bias correction for six indices generated from the three variables of bottom temperature, sea ice, and net primary production.

Bias correction uncertainty was a large source of uncertainty for the indices aside from bottom temperature, particularly for those that are associated with seasonal cycles and capture extrema in time series or phenology (Figure 7). For example, both bloom start date and ice-free period had greater than 25% of total uncertainty stemming from bias correction uncertainty throughout the duration of the time series. Net primary productivity, fraction ice coverage, and monthly-averaged September net primary production all showed around 10-25% of uncertainty being attributable to bias-correction approaches for the duration of the time series. For bottom temperature, uncertainty due to bias correction was low compared to the other five indices, but like fraction ice coverage and the number of ice-free weeks, scenario uncertainty became a large source towards the end of the century (Figure 7).

#### Case studies

##### I. Forecasting Pacific cod thermal spawning habitat

Predicted thermal spawning habitat suitability for Pacific cod in the eastern Bering Sea differed depending on the bias correction method applied to the bottom temperature time series, as well as the simulation (i.e., which parent model and SSP; Figure 8). Across the entire time series, for a given parent model and emission scenario combination (e.g., low emission CESM simulation), the largest percent difference between thermal spawning habitat suitability calculated from the nine different time series of temperature (one uncorrected + eight bias corrected) in a given year ranged from 3.9% for the low emission CESM simulation to 149.3% for the high emission MIROC simulation. In other words, the estimated thermal spawning habitat suitability index for a given year varied by at least 3% and up to ∼150%. The largest differences among thermal spawning habitat suitability from different bias-correction approaches were evident from mid- to late-century and for the high emission scenario of the CESM and MIROC simulation (Figure 8).

**Figure 8.**
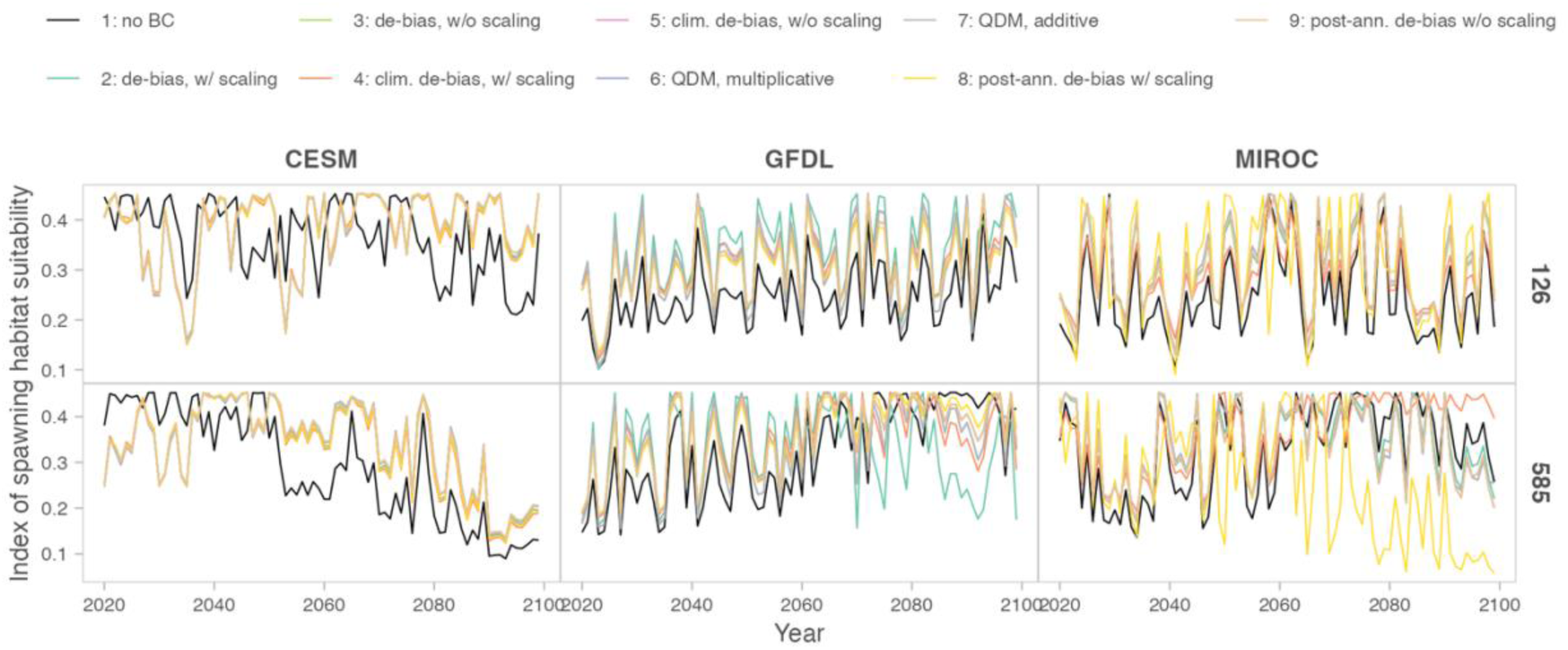
Annual time series of the index of spawning habitat suitability for each bias correction method (colors) and simulation (columns/rows).

For all simulations, thermal habitat suitability ranged from ∼0.5 to 0.45 (Figure 8). Thermal habitat suitability was highly variable regardless of the bias correction approach applied to bottom temperature (Figure 8). The habitat suitability time series generated from CESM differed from those from the other two parent models, likely due to the differences in temperature across the parent models (CESM is the warmest, Figure 4). For all bias correction approaches, the thermal habitat suitability time series from the low emission scenario of CESM bottom temperature was variable with no trend, while the thermal habitat suitability time series from the high emission scenario appeared to decrease over time, starting around mid-century (Figure 8). For CESM, the raw (no bias correction) time series of temperature predicted lower spawning habitat suitability per year than the time series calibrated with bias-correction methods. The thermal habitat suitability time series generated from the different bias correction approaches for the GFDL and MIROC bottom temperature time series were generally all in agreement, particularly for the time series under the low emission scenario. For the high emission scenario, some methods predicted higher or lower thermal habitat suitability than other bias correction approaches (Figure 8). For example, the bias removal and rescaling method (#2), predicted lower thermal habitat suitability starting around 2070 under the high emission scenario of the GFDL model, and the post-annual bias removal and rescaling method (#8) predicted lower thermal habitat suitability. While highly variable, most time series (all parent models, emission scenarios, and bias-correction method, n = 54) exhibited no real long-term increasing or decreasing trend from 2020 - 2099.

##### II. Sea ice & zooplankton

Predicted zooplankton abundance differed considerably across time depending on the bias-correction method applied to the sea ice time series (Figure 9). These differences did not necessarily mirror those observed for the Pacific cod thermal spawning habitat suitability predictions. For example, differences in zooplankton abundance predicted from sea ice adjusted with the eight bias-correction methods and the raw time series differed in terms of general patterns among simulations and emission scenarios. The largest percent difference between yearly predictions of zooplankton abundance across the nine time series ranged from 12.7% for the low emission GFDL simulation to 151.5% for the high emission MIROC simulation. Although the minimum percent difference between the time series was greater than that of the Pacific cod thermal spawning habitat suitability case study, the highest percent differences were similar and were both stemming from the high emission MIROC simulation.

**Figure 9.**
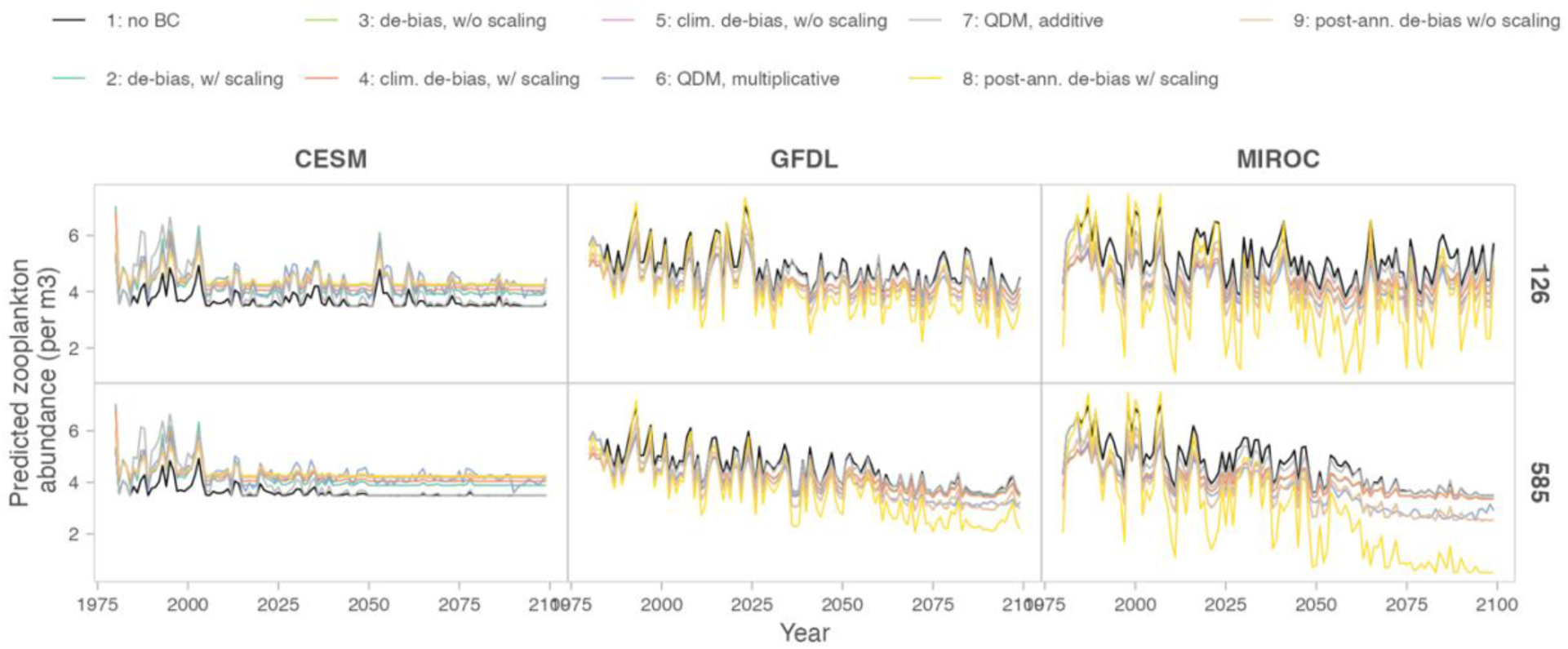
Predicted zooplankton abundance based on projected sea ice time series bias corrected in eight different ways (colors), as well as the raw time series (black), for each parent model (column) and emission scenario (rows).

Although long-term trends of zooplankton abundance were similar across all time series with different bias correction approaches applied, the magnitude differed (Figure 9). There was a greater difference in magnitude across time series under the high emission scenarios for both GFDL and MIROC, whereas the magnitude of difference among time series was similar for both emission scenarios for CESM (Figure 9). Predictions of zooplankton based on time series of sea ice from the CESM model for all bias correction methods differed from both GFDL and MIROC (Figure 9), likely due to the differences in sea ice prediction (CESM is the warmest; Figure 4). Under both high and low emission scenarios of CESM, the raw (#1) and the additive quantile delta mapping (#7) time series of predicted zooplankton abundance were the lowest, but under both emission scenarios of GFDL and MIROC, these same two time series were the highest. For CESM, the remaining seven time series of zooplankton abundance predictions were generally similar, with less variability in the annual bias correction approaches (#8 and #9). For the high emission scenario of GFDL and MIROC, there were three groupings, similar as described above for the Pacific cod thermal spawning habitat suitability index. In terms of magnitude, the raw (#1), the additive quantile delta mapping (#7), bias removal with and without rescaling (#2 and #3), and the climatological bias removal with and without rescaling (#4 and #5) predicted the highest zooplankton abundance (Figure 9). The next lowest were multiplicative quantile delta mapping (#6) and the post-annual bias removal without scaling (#9). Finally, the post-annual bias removal with rescaling predicted the lowest zooplankton abundance (Figure 9).

##### III. Phytoplankton and zooplankton phenology

The degree of match-mismatch between the phytoplankton and zooplankton peak blooms over time depended on the bias correction method applied, as well as the parent model and emission scenario (Figure 10, Table S1). For most bias correction methods, parent models, and emission scenarios, the number of days between the spring phytoplankton and zooplankton blooms (match-mismatch) did not differ considerably over time. This was true for 47 of the 54 total time series (nine bias correction approaches [eight bias corrected, one raw] x three parent models x two emission scenarios). However, for seven of the time series, there was a significant trend such that over time, there was a greater match in peak bloom date over time (the number of days between the peak of the phytoplankton and peak of the zooplankton bloom decreased) (Table S1, Figure 10). Four of these time series applied a climatological bias removal with rescaling correction (#4), two applied a climatological bias removal without rescaling correction (#5), and the last one applied a post-annual bias removal without rescaling (#9) approach. Of the seven time series where there was a long-term trend in the mean number of days between blooms, the parent models and emission scenarios differed (Table S1, Figure 10). Four time series were CESM (high and low emission scenarios using the climatological bias removal with and without rescaling methods, #4 and #5), two were MIROC (high emission scenario for climatological bias removal without rescaling [#5] and post-annual bias removal without rescaling [#8]), and one as GFDL (high emission scenario of climatological bias removal without rescaling [#5]).

**Figure 10.**
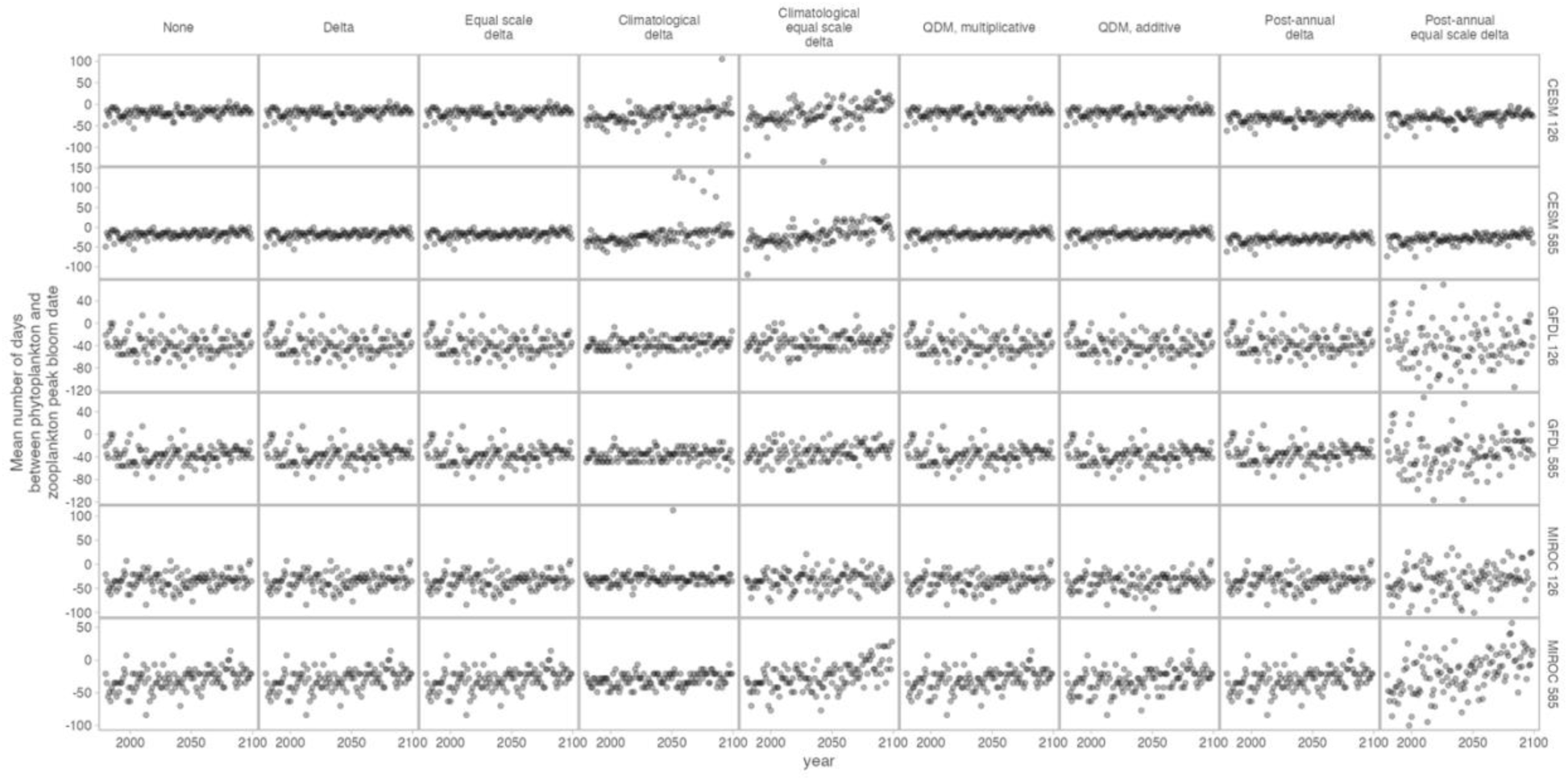
The number of days (match-mismatch) between the peak day of year of the phytoplankton spring bloom and the peak day of year of the zooplankton spring bloom depends on bias correction method applied (columns) and simulation (rows).

The variability in the mean number of days between the phytoplankton and zooplankton spring blooms for a given year varied greatly across the different time series associated with the different bias correction approaches applied, parent models, and emission scenarios (Figure 10). There was very little variability over time in match-mismatch for both emission scenarios of CESM, with the exception of time series generated from both climatological bias correction approaches (climatological bias removal with and without rescaling, #4 and #5). There was more variability in the time series of other parent models; for example, match-mismatch for a given year was more variable for both emission scenarios of GFDL and MIROC compared to both emission scenarios of CESM. Time series of match-mismatch generated from post-annual bias removal without rescaling correction (#9) were the most variable out of any other bias correction approach, followed by both climatological bias correction methods (#4 and #5; which largely suggested the change in match-mismatch over time, see above). In contrast, the time series of match-mismatch generated from the climatological bias correction approaches (#4 and #5) for the simulations that did not show a trend were actually less variable than when other bias correction approaches were applied (Figure 10).

## 4 Discussion

Overall, we found that the choice of bias correction method, and the time scale over which it is applied, contribute significantly to the uncertainty surrounding future projections and the biological models forced with them. All three variables - bottom temperature, net primary productivity, and sea ice - had inherent biases, necessitating some form of bias correction. The trajectory of a given variable differed (sometimes considerably) across bias correction methods, with some methods even amplifying biases. The amount of uncertainty attributable to internal variability, the suite of parent models and emission scenarios, and bias correction varied across the six indices generated from the three variables: annually-averaged bottom temperature, annually-averaged net primary production, spring bloom start date, monthly-averaged September net primary production, maximum fraction ice coverage, and number of ice-free weeks per year. Bias correction uncertainty was an important source of uncertainty for all variables and contributed considerably to total uncertainty for seasonal variables. Differences across projections were reflected in the biological models, where predictions of Pacific cod thermal spawning habitat suitability, zooplankton abundance, and phenology of lower trophic levels differed considerably depending on which bias correction method was applied to the time series that forced the model.

Of the bias techniques considered in this paper, the multiplicative quantile delta mapping method (#6) was the only method that maintained numerical realism across all variables at the original weekly temporal resolution. Quantile mapping, a category of bias correction methods that shifts values such that the resulting cumulative distribution functions (CDF) are in line with that of the observations, is among the most widely used type of approach (Piani et al. 2010, Cannon et al. 2015, Dinh and Aires 2023). In addition to correcting the CDF to match that of the observations, this approach additionally preserves the change signals of the model in terms of quantiles (Cannon et al. 2015, see 2.2.3 Bias correction methods section). Other techniques we explored produced a number of artifacts. For example, the majority of methods produced values outside the range of values physically possible (e.g., net primary productivity rates less than zero, ice fraction less than zero or greater than one), especially when confronted with variables with strong seasonal variations in bias. Several techniques also produced undesirable features that disrupted the expected seasonal cycle when confronted with phenological biases (e.g., climatological bias removal with and without rescaling, #4 and #5). Such findings, e.g., differences across time series depending on bias correction method applied, undesirable artifacts, etc., were also found in other studies comparing the effect of bias correction techniques on projections from climate models (Watanabe et al. 2012, Dinh & Aires 2023). However, most of these studies and those that apply quantile mapping methods are focused on terrestrial applications and fewer indices (e.g., temperature or precipitation; Piani et al. 2010, Watanabe et al. 2012, Cannon et al. 2015). Here, we found the multiplicative quantile delta mapping approach to also be ideal for marine applications.

The results of our uncertainty source partitioning showed many of the same characteristics as previous work (Frölicher et al. 2016). In Frölicher et al. (2016), variables tightly coupled to physical dynamics, such as temperature and pH, were dominated by scenario uncertainty, while more biologically-influenced variables like net primary productivity and low-oxygen waters were instead dominated by model uncertainty. We observed similar patterns, with net primary productivity, bloom start date, and September net primary production dominated by model, internal variability, and bias correction method uncertainties. Model uncertainty remained a high contributor across our suite even though all regional models were run with identical biological dynamics, similar to other models and variables for both marine and terrestrial global and regional ESMs (Frölicher et al. 2016, Bonan and Doney 2018, Lehner et al. 2020). In our case, this confirms that it is not simply differences in the structure of the biological equations within parent models that could lead to a spread across projections, but rather the nonlinear manner in which biological models propagate physical signals from the ocean into ecosystem metrics. Given this relationship, it is not surprising that variables that indicate high sensitivity to parent model uncertainties are also those most strongly influenced by the choice of bias correction method. Within our suite of example indices, bias correction added only a small amount of uncertainty to bottom temperature, a variable where the biases seen across parent models was small relative to the long-term projected change over the 21^st^ century. However, the uncertainty was much higher around the other variables (i.e., those in Figure 7), with bias correction technique uncertainty contributing equally or surpassing the amounts of uncertainty derived from more commonly-considered sources like interannual variability, model structural uncertainty, and scenario uncertainty. This can also be seen in our ice-related indices, which are both tightly coupled to temperature (and temperature biases), and which are numerically limited to a small range. In particular, the ice-free period index combines several features that make bias-correction particularly complex: it is based on a numerically-constrained variable with seasonally-varying biases and attempts to quantify a tipping-point when the system crosses a fixed threshold. By the end of century, the uncertainty in this variable was 36.4 weeks, 70% of the maximum theoretically possible uncertainty (52 weeks) for this quantity. This results in opposite projections highlighting the complexity and importance of bias correction: the Bering Sea may be prone to completely ice-free years in the near-future and permanently ice-free beginning around 2060, or, alternatively, seasonal ice may remain a stable feature of the Bering Sea well past the end of the century.

The differences in bottom temperature, sea ice, and net primary production due to bias correction method applied were reflected in the downstream biological models forced with these variables. These differences ranged from differences in magnitude, variability, and trajectory of Pacific cod thermal spawning habitat suitability, zooplankton abundance, and phenology of lower trophic levels. For example, the magnitude and variability of expected Pacific cod thermal spawning habitat suitability to the end of century greatly differed depending on which bias correction method was applied, whereas in this case, differences in trajectory were largely due to differences in bottom temperature projections among parent models. For the phenology of lower trophic levels, the number of days between the peak date of the phytoplankton bloom and peak date of zooplankton abundance were either getting smaller (peaks closer together) or not changing, depending on the bias correction method applied. The variability in the number of days between peaks also varied considerably depending on the bias correction method applied. The trajectories and variability of predicted zooplankton abundance were largely similar over time regardless of the bias correction approach applied to the sea ice that forced the model. However, the magnitude of abundance greatly varied depending on which bias correction method was applied to the sea ice time series. Collectively, we conclude that choice of bias correction method affects our understanding of how biology may change in the future with climate change. While other studies on bias correction approaches are largely focused on terrestrial models and do not assess the effect of bias correction method further downstream (i.e., in a model that uses the bias-corrected variable as done in this study; Frölicher et al. 2016, Lehner et al. 2020, Pozo Buil et al. 2023), it is clear that guidance is needed to determine how to approach bias correction for use in downstream biological models.

We noted earlier (section 2.2.1) that the motivation for bias correction is usually tied to difficulty of interpreting unadjusted ESM output in a biological context. However, as demonstrated in our examples, bias correction does not entirely solve this issue. While numerical techniques can make the data more convenient to work with, that does not necessarily eliminate the ambiguities that surround interpretation of either biased or bias-corrected ESM output. For example, if a model predicts loss of habitat from a location that should not have habitat in the first place, statistical manipulation can only go so far to aid in how one should interpret that predicted loss. If a model underestimates present day habitat and therefore underpredicts the end-of-century change due to hitting a hard threshold, statistical correction cannot answer whether the dynamics would have led to a larger change had the threshold been further away. Full interpretation of ESM output requires a nuanced understanding of why the biases may have arisen in the underlying ocean model and whether useful information can be gleaned from the model (with or without bias correction) despite that. Indeed, the choice of bias correction method depends on application and what should be emphasized from the observations - average mean, high resolution pattern - and on the model data - trend, distribution of changes (Dinh & Aires 2023, Ho et al. 2012, Teutschbein and Seibert 2012).

Our study highlights that there is no one-size-fits-all technique that conveniently addresses all types of biases across all variables. Yet, we recognize the need to provide guidance on which bias correction approach to use and how to select a bias correction approach. Given that quantile delta mapping could be successfully applied “out of the box” to a variety of variables with widely-differing underlying distributions and biases, the QDM technique may be well-suited to be applied to time series of climate model variables that force downstream biological models. However, we also note that this is one of the more complex techniques to implement (Cannon et al. 2015). While certainly not a difficult calculation, the need to calculate empirical CDFs, extract inverse values from them, apply sliding windows, etc., offers more room for small variations in the precise implementation of the calculation across end users and their preferred high-level modeling languages than techniques centered simple mean and variance mapping as in the bias removal methods (Cannon et al. 2015, Ho et al. 2012). The practical advantages of consistent application of a simple bias correction technique for all users of an ocean model (e.g., Bering 10K) may outweigh the numerical advantages of a more complex technique. We also note that many users of the Bering10K output in the ACLIM project applied a log transformation prior to bias correction for variables with lognormal distributions (as opposed to being normally distributed). After log transforming, these time series were then bias corrected and then back-transformed. Like the constraints of the bias removal with and without rescaling methods (#1 and #2), this approach would only work if the variable was truly lognormally distributed. Additionally, this resulted in extreme and unrealistic values and so we recommend the CDF-based methods (QDM) as they are robust to a variety of distributions. To that end, we recommend as much transparency as possible when applying any form of bias correction such that end users can make an informed decision as to whether the technique is appropriate for their research use case. We also suggest that analysts assess the effect of different bias correction methods on projections as done here. To assist with navigating the bias correction process for ESM output, we present a decision tree (Figure 11). Based on our work here, this tree outlines the steps analysts could take to navigate the entire bias correction process - from the raw ESM model output to assessing the effect of bias correction methods.

**Figure 11.**
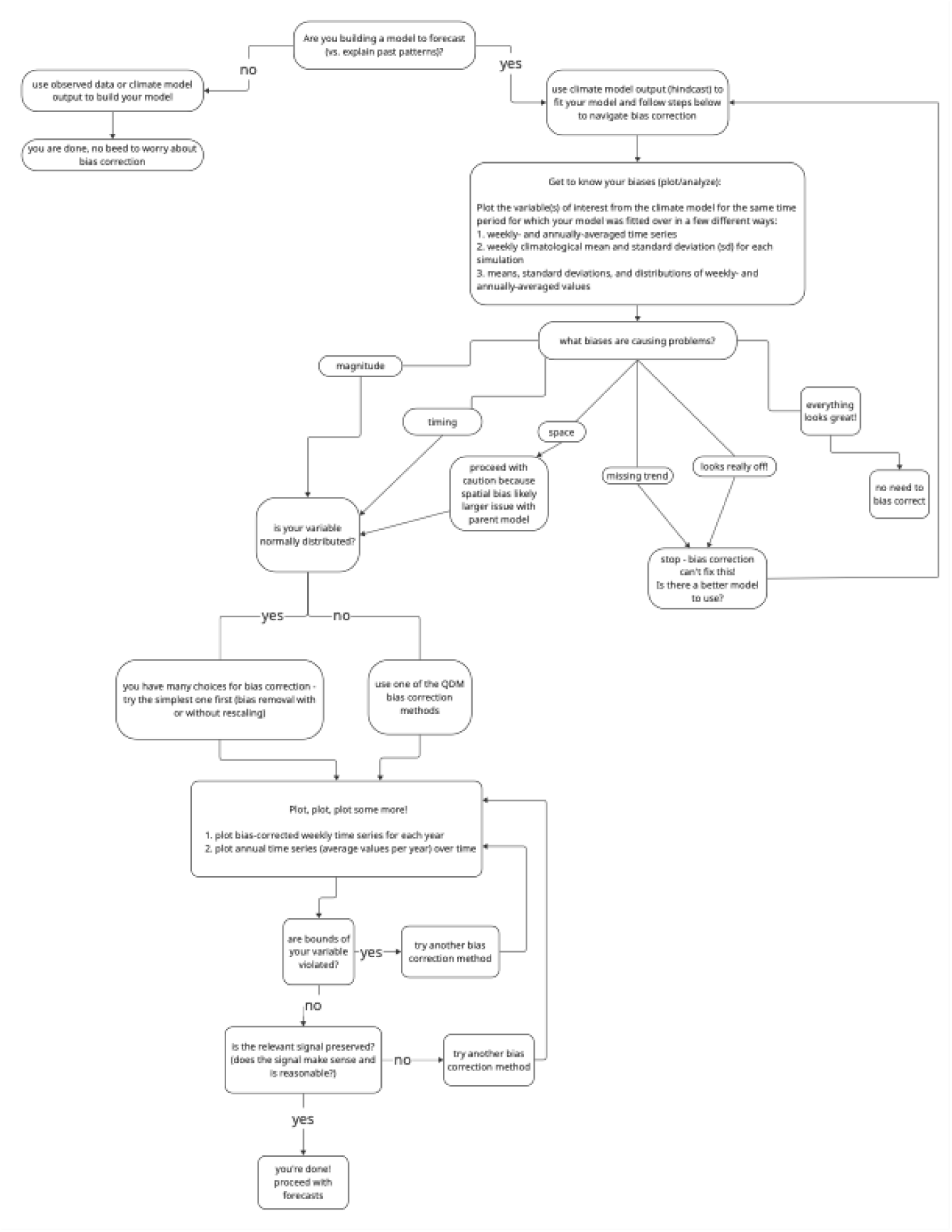
A decision tree for analysts using time series from global or regionally-downscaled Earth System Models (ESMs) to understand the physical changes in an ecosystem(s) over time or to use the physical variables as inputs in statistical models to understand change in biological systems over time.

There are still remaining avenues for identifying biases in ESM output and applying bias correction techniques to ameliorate these biases. First, multivariate bias correction, or bias correcting multiple variables so as to not disrupt the dependency structure between or among variables is another option to consider (Vrac and Friederichs 2015, Maraun 2016, Vrac 2018). However, using a multivariate bias correction approach is not trivial and most examples in the literature only consider two variables at a time (e.g., Vrac and Friederichs 2015). Bias correction pre-downscaling would avoid the need to bias correct post-downscaling and forestall multivariate bias correction, yet not doing so may be beneficial (Dinh and Aires 2023, Pozo Buil et al. 2023). For example, if a model is bias corrected prior to downscaling, the spatial patterns in the past are projected into the future (Dinh and Aires 2023, Pozo Buil et al. 2023). Second, the spatial scale at which bias correction occurs could be considered. One could bias correct values for every grid cell individually, or aggregate over space. For example, Bigman et al. (2023) used the bias removal with rescaling method (#2) to bias correct bottom temperature, but the factors that comprise the scaling factor in the equation (the standard deviation of the hindcast during a reference period divided by the standard deviation of the raw projections over the same reference period) were averaged by a certain spatial structure. While this did not affect the results in that study, the spatial scale of each parameter in the equation could be considered (Bigman et al. 2023). Finally, the choice of datasets differ across methods (i.e., observations, hindcast, and ‘historical’ time series [the historical portion of the forecast simulation]) and it is likely that the result of bias correction will differ depending on the dataset used in different parts of the process.

Here, it was our goal to identify how various established bias correction methods affected our understanding of how environmental conditions may change in the future, and how variation due to different methodologies may be propagated to downstream ecological models. The latter step is an important advancement of previous comparisons of different bias correction methods, which have largely focused on the output of ESMs rather than downstream models. Through our effort, this work highlights the complexities of bias correction, makes suggestions for how to deal with these complexities and uncertainties, and outlines a step-by-step decision tree to identify the best bias correction method for many variables from regional and global ESMs. Indeed, our findings are broadly applicable to other ESMs. First, the decision tree is applicable to any analyst using uncorrected ESM output (i.e., from a model that has had no bias correction applied). Second, we assessed the effect of bias correction on many different variables that could be extracted from an ESM, including the most commonly used methods and those that represent different scales and biases (e.g., timing of seasonal events). Forecasting the effects of climate change in biological systems is complex, challenging, and uncertain. Our work aims to help reduce the uncertainty in our understanding of future physical and biological changes.

## Acknowledgements

We thank the EcoFOCI group at NOAA Alaska Fisheries Science Center for data and those who have conducted surveys to collect and process these data. We also thank the Plankton Sorting and Identification Center in Szczecin, Poland for zooplankton identification. We thank Lauren Rogers, Dave Kimmel, and Jens Nielsen for advice and input on this manuscript. We also thank the CEFI and Cookies team and the wider ACLIM group for their comments and suggestions, and those who developed and implemented the Bering 10K model.

## Author Contributions

JSB: conceptualization, data curation, formal analysis, methodology, project administration, software, visualization, writing – original draft, writing – review and editing; KAK: conceptualization, data curation, formal analysis, methodology, software, visualization, writing – original draft, writing – review and editing; KKH: conceptualization, data curation, methodology, writing – review and editing.

## Supplement

**Table S1.**
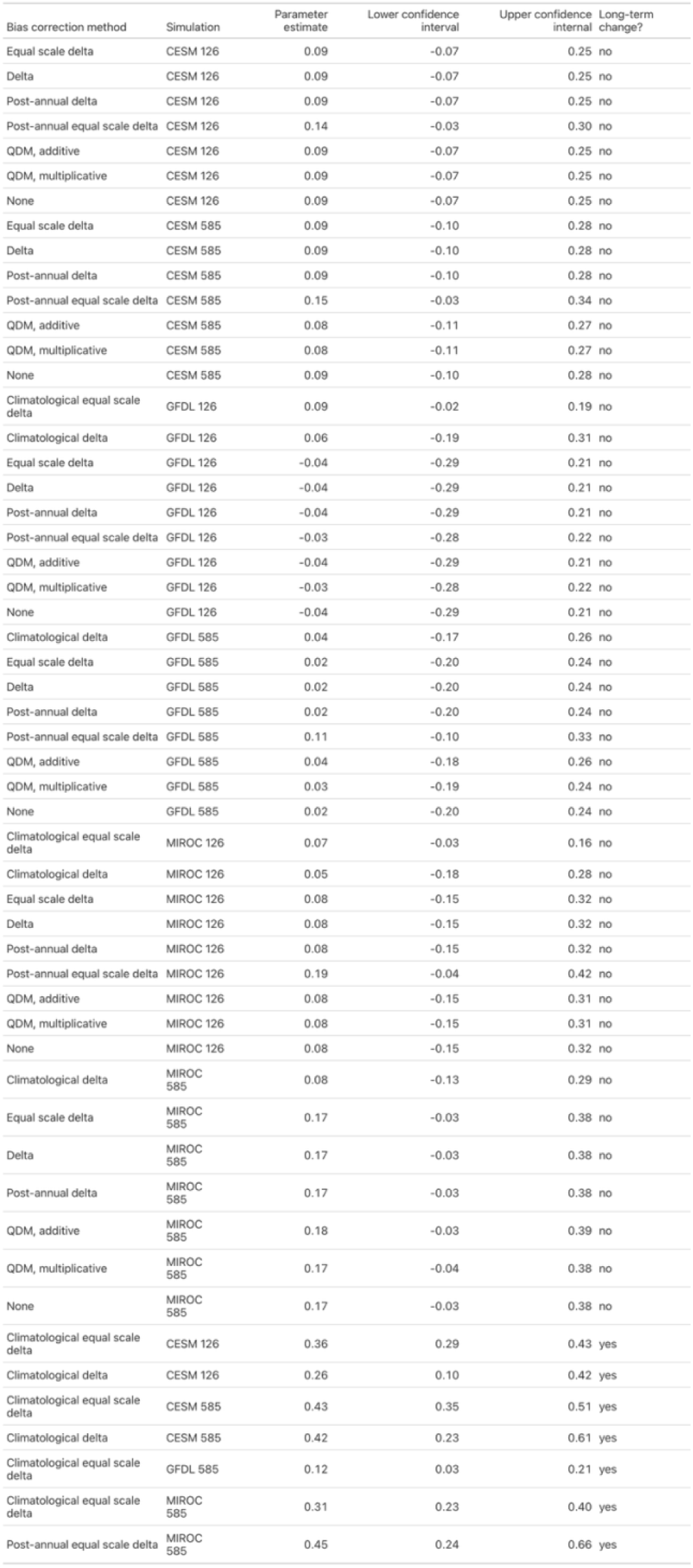
Regression coefficients for linear model fits to assess the presence of a long-term change in the match-mismatch between the phytoplankton and zooplankton spring blooms.

**Figure S1.**
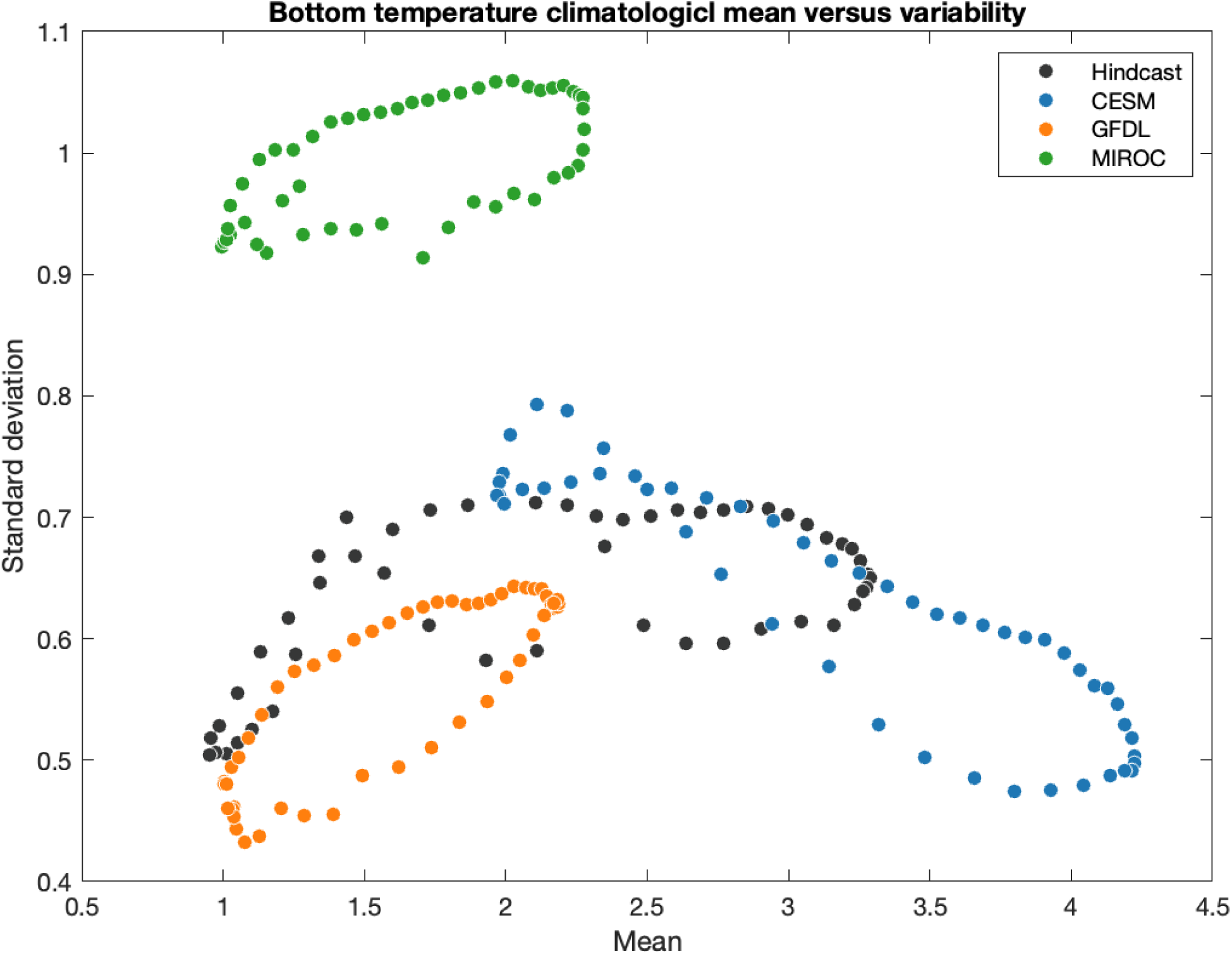
Climatological mean versus standard deviation across each model reference period.

