## Supplementary Material for "Bias correction for integrated climate projection modeling and its effect on downstream biological models"

Table S1. Regression coefficients for linear model fits to assess the presence of a long-term change in the match-mismatch between the phytoplankton and zooplankton spring blooms.

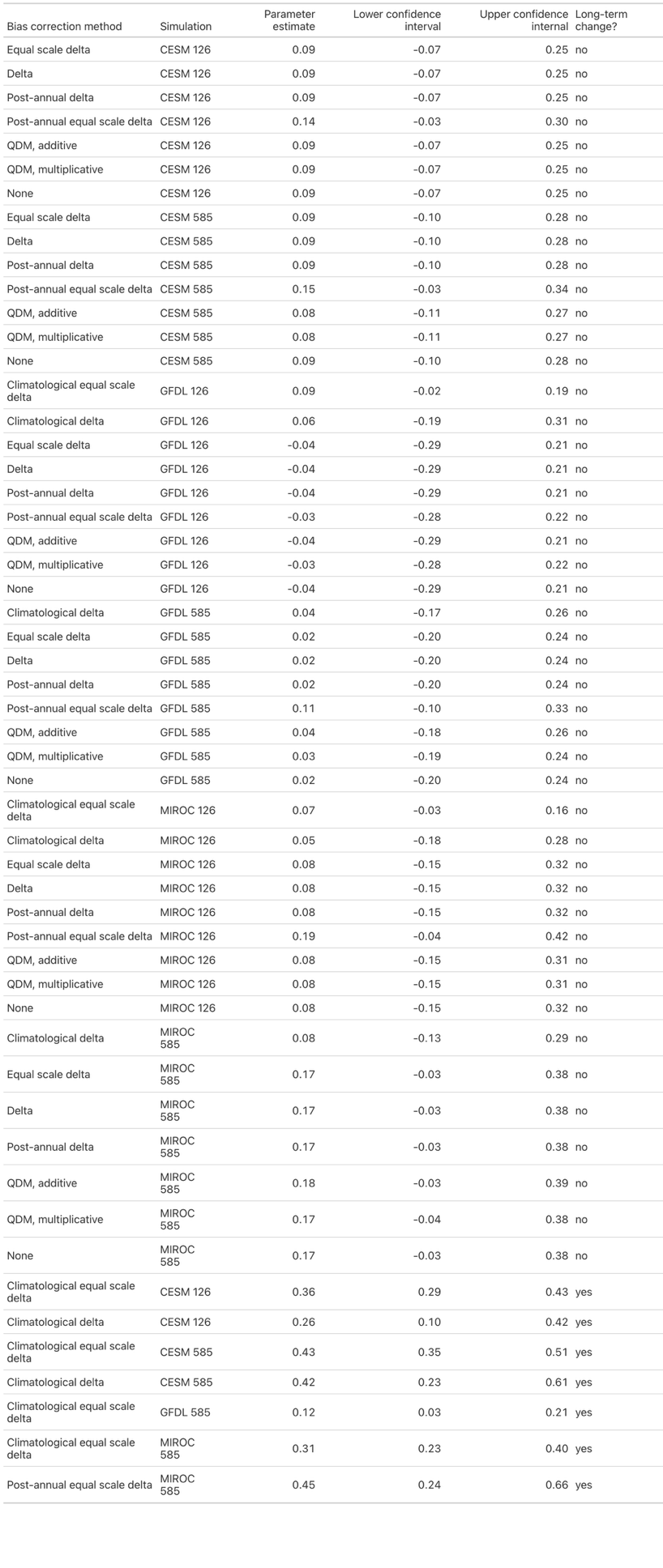

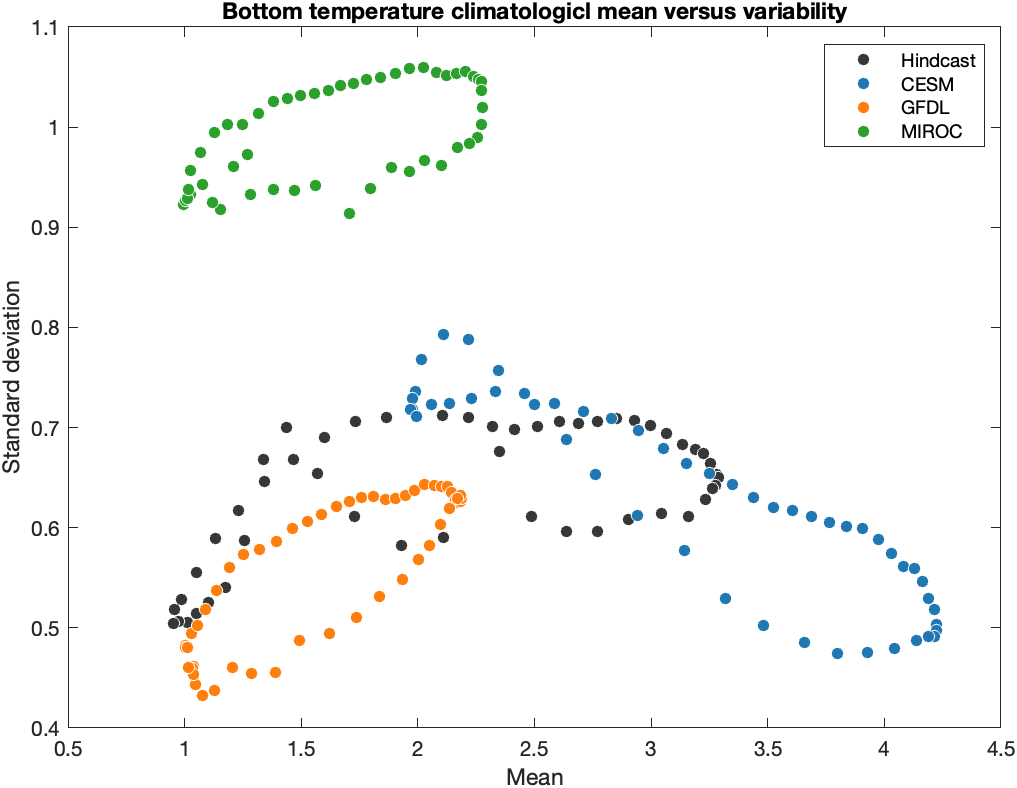

Figure S1. Climatological mean versus standard deviation across each model reference period.
